# Transcriptional units form the elementary constraining building blocks of the bacterial chromosome

**DOI:** 10.1101/2022.09.16.507559

**Authors:** Amaury Bignaud, Charlotte Cockram, Eric Allemand, Julien Mozziconnacci, Olivier Espeli, Romain Koszul

**Affiliations:** Institut Pasteur, CNRS UMR 3525, Université Paris Cité, Unité Régulation Spatiale des Génomes, F-75015 Paris, France; Center for Interdisciplinary Research in Biology (CIRB), Collège de France, CNRS, INSERM, Université PSL, Paris, France; Muséum National d’Histoire Naturelle, Structure et Instabilité des Génomes, UMR 7196, Paris 75231, France; INSERM - U1163, Unité mécanismes cellulaires et moléculaires des désordres hématologiques et implications thérapeutiques, Institut Imagine, Paris, France; Collège Doctoral, Sorbonne Université, F-75005 Paris, France

## Abstract

Transcription generates local topological and mechanical constraints along the DNA fiber, driving for instance the generation of supercoiled chromosomal domains in bacteria. However, the global impact of transcription-based regulation of chromosome organization remains elusive. Notably, the scale of genes and operons in bacteria remains well below the resolution of chromosomal contact maps generated using Hi-C (~ 5 – 10 kb), preventing to resolve the impact of transcription on genomic organization at the fine-scale. Here, we combined sub-kb Hi-C contact maps and chromosome engineering to visualize individual transcriptional units (TUs) while turning off transcription across the rest of the genome. We show that each TU forms a discrete, transcription-induced 3D domain (TIDs). These local structures impose mechanical and topological constraints on their neighboring sequences at larger scales, bringing them closer together and restricting their dynamics. These results show that the primary building blocks of bacteria chromosome folding consists of transcriptional domains that together shape the global genome structure.

## Introduction

Bacterial genomes are organized into the nucleoid, a well-defined physical object where DNA, RNA and proteins interacts (Kleckner et al. 2014; Lioy et al. 2021). These interactions locally shape the conformation of the Mb long circular chromosome (Dame et al. 2020). DNA helicases, involved for instance in replication and transcription, modulate transiently the supercoiling level of the DNA fiber (Liu and Wang 1987) by creating twin-domains spanning 25kb in each direction (Visser et al. 2022). Topoisomerases, mainly Topo I and DNA gyrase, maintain supercoiling homeostasis, to keep the negatively supercoiled state necessary for DNA compaction and strand opening operations (Dorman 2019). Radial plectoneme loops are proposed to decorate the chromosome, either in association with protein complexes of the structural maintenance of chromosome (SMC) family (Mäkelä and Sherratt 2020; Ganji et al. 2018) or with supercoil-induced processes (Deng et al. 2004; Postow et al. 2004). Bacterial chromosomes Hi-C contact maps have also revealed higher-order levels of organization (Le et al. 2013; Le and Laub 2016; Lioy et al. 2018; Marbouty et al. 2015; Umbarger et al. 2011), with directionality index analysis (a statistical parameter that assesses the degree of upstream or downstream contact bias for a genomic region) pointing at ~30 chromosome self-interacting domains (or CIDs) ranging in size from ~30 to 300 kb (Le et al. 2013). A careful analysis further unveiled a correlation between highly expressed, long genes (HEGs) and CID boundaries, though not systematic (Marbouty et al. 2015; Le and Laub 2016), while a general genomewide correlation was further described between transcription level and contact frequencies between pairs of adjacent, 5 kb DNA segments (bins) (Lioy et al. 2018). Furthermore, inhibition of transcription initiation by rifampicin abrogates domains and decondense nucleoids within minutes, suggesting a direct role for transcription in folding the chromosome (Worcel and Burgi; Stracy et al. 2015). In agreement with the idea that transcription shapes chromosome organization, recent experiments and biophysical models revealed that RNA production reduces the solvent quality of the cytoplasm for the chromosome and consequently impacts its local conformation (Xiang et al. 2021). However, with respect to the scale of gene and operon (<10 kb) (Deng et al. 2004) of bacterial genome, these analyses remain relatively coarse. In addition, gene density, concomitant transcription, and cell-to-cell variability of hundreds of genes could lead to intermingled patterns, leaving the possibility that fundamental underlying structural features have been overlooked.

Here, we combine a high-resolution Hi-C protocol recently adapted for bacteria (Cockram et al. 2021a) with chromosome engineering and cellular imaging to address the link between chromosome architecture and transcription at a higher level. We show that all active TUs form discrete individual 3D domains that form the primary building blocks for larger chromosome folding.

## Results

### High-resolution contact maps reveal transcription associated local chromosome interactions

The 1 kb resolution Hi-C contact map of exponentially growing *E. coli* cells (methods) unveil a strong heterogeneity in the short-range contact signal (**Figure 1a**), with ~200 denser, thick bundles along the main diagonal (see Methods for bundle calling). These patterns, which cover approximately 1,300 kb, are strongly correlated with transcriptional activity and disappear upon addition of rifampicin (**Figure 1b**; Supplementary Data Figure 1a, b). They range in size from 1 to 20 kb and are distributed over the entire genome map (Supplementary Data Figure 1c). The potential to make protein-DNA crosslinks will influence local Hi-C contacts (Scolari et al. 2018). Therefore, the local protein concentration on the DNA (protein occupancy) may contribute to the local Hi-C bundle signal. We took advantage of recent high resolution maps of protein occupancy on the *E. coli* genome (Freddolino et al. 2021) to test whether silent regions nevertheless strongly enriched in proteins (EPODs) would appear as local bundles in Hi-C maps. As shown on **Figure 1c**, only ~10% of EPODs regions appear involved in a bundle, suggesting that protein occupancy per se is not sufficient to promote their formation of bundle domains.

**Figure 1:**
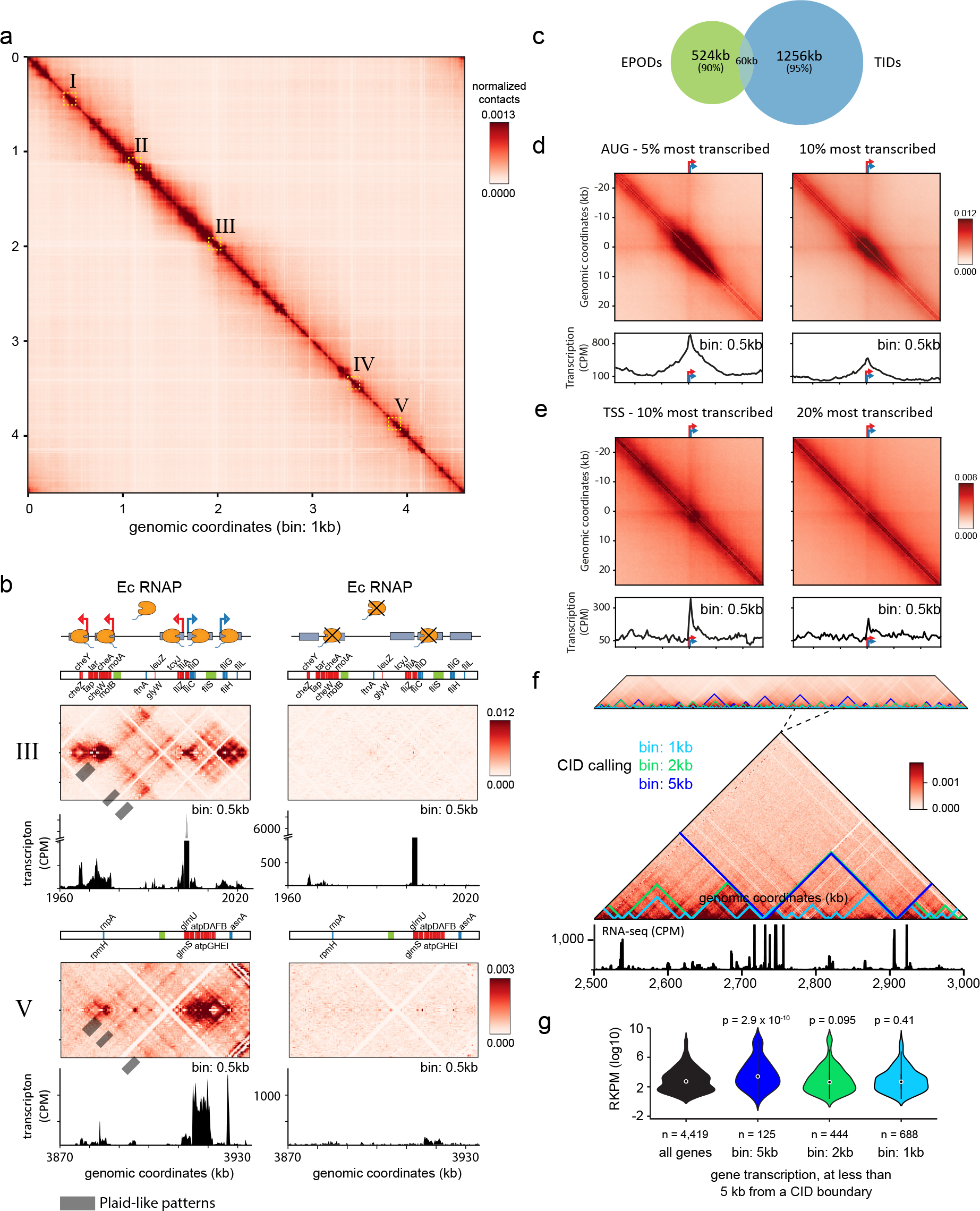
The bacterial chromosome is structured by tens of small transcriptionally active 3D units. **a,** Hi-C normalized contact map of wt *Escherichia coli* cells (bin: 1kb). The five yellow squares I – V underline representative 64 kb regions magnified in either panel **b**) or in **supplementary Figure 1**. **b,** Magnifications of regions III and V in absence (left) and presence (right) of rifampicin. Ec RNAP: *E. coli* RNA polymerase II. For each window and condition, a schematic representation of the region's genetic content is presented on the top, with the names of genes within the 10% most transcribed indicated in blue and red for forwards and reverse orientation, respectively and silent EPODs regions (green) (Freddolino et al. 2021). Middle panel: normalized contact map (bin: 0.5 kb). Lower panel: RNA-seq profile in Count Per Million (CPM). **c,** Venn diagram of EPODs labelled regions (Freddolino et al. 2021) and of regions labelled as thick bundles. The metric used correspond to the total size of the corresponding regions, in kb. **d,** Top: pileup of contact map 50 kb windows (bin: 0.5 kb) centered on the start codons (AUG) of the 5% (left) and 10% (right) most transcribed genes of the genome. Bottom: corresponding pileup of transcription (RNA-seq) tracks. **e,** Pileup of contact map 50 kb windows centered on the transcription start sites (TSS) of the to 10% and 20% most transcribed TUs (i.e. operons). Bottom: corresponding pileup of transcription (RNA-seq) tracks. **f,** Self-interacting domains called using DI analysis on contact maps binned either at 1 kb (cyan), 2 kb (green) and 5 kb (blue). Top, visualization of domains over the entire genome. Middle, magnification of a 500 kb region. Below, Corresponding RNA-seq track in CPM. **g,** Violin plot distributions of transcript levels for all genes in the genome (black), and for all genes in 10 kb windows centered on the domain boundaries called on the 5 kb (blue), 2 kb (green) and 1 kb (cyan) bin maps. The p-values of non-parametric Mann-Withneyu test of whether the later distributions follow a genomewide distribution are indicated.

In addition, a plaid-like pattern was often observed, corresponding to enrichment in contacts between successive transcribed DNA regions, alternating with non-transcribed region with which they make fewer contacts (i.e. resulting in “empty” stripes) (**Figure 1b**; Supplementary Data Figure 1a). The positioning of the “empty” stripes of the plaid patterns are not correlated with GC%, restriction site density, nor protein occupancy (as quantified in (Freddolino et al. 2021)) suggesting that they do not correspond to DNA regions that are poorly visible and/or less captured by the Hi-C protocol (Supplementary Data Figure 1a, b; Methods) (Cournac et al. 2012). This observation suggests that transcribed regions tend to contact each other locally, either because they tend to relocate to the nucleoid external periphery, as suggested by super resolution imaging (Stracy et al. 2015; Gaal et al. 2016), or through transcription-mediated clustering. To quantify the correlation between contacts and gene expression, a pile-up analysis of contacts centered on the start codon of the 5 and 10% most transcribed genes was performed (**Figure 1c**). A large contact signal centered on the start codons appeared, strongly correlated with the corresponding averaged transcription signal (Pearson correlation: 0.81). Because bacteria genes are often organized into operons and cotranscribed, we then plotted the pile-up contact windows centered on the start codon of the first gene of the most transcribed operons (TSS) (**Figure 1d;** Methods). The pile-up displays an enrichment in contact signal that increases abruptly precisely at TSS positions, and extends over the area spanned by the transcription track, further reinforcing the notion that short-range (0-5kb) Hi-C contacts are correlated with transcription levels (Pearson correlation: 0.62). A slight enrichment of contacts between the TU and upstream and downstream regions is also observed, a signal that corroborates the plaid-like pattern observed on the sub-kb contact map.

Taken together, these observations suggest that the primary blocks organizing the *E. coli* chromosome are a succession of transcriptionally induced domains, or TIDs, that appears as thick bundle in Hi-C contact maps and can interact together as long as the genomic distance between them remains small (<25 kb).

### Transcription Induced Domains explains CIDs detection in low-resolution maps

We next compared the positions of the TUs with the CIDs boundaries previously identified along the *E. coli* genome (Methods). CIDs called as previously done (Lioy et al. 2018) in the contact map binned at 5 kb revealed 27 domains. 22 boundaries overlapped those previously identified, while the differences remain just at the edge of the detection threshold (blue signal, **Figure 1e**; Supplementary Data Figure 1c; Supplementary data Table 1). As shown, these boundaries are enriched with HEGs (**Figure 1f**; Supplementary Data Figure 1d). The same DI analysis on a 2 kb binned map yielded 30 new boundaries (green signal, **Figure 1e,f**; Supplementary data Table 1), but proved too noisy when applied on a 1kb matrix (Supplementary Data Figure 1e, Supplementary data Table 1) (as the width of the vector used relies on the resolution). To detect CID-like signals in 1kb contact maps, we adapted HiC-DB, another insulation score approach (Chen et al. 2018) (Methods). We detected 135 boundaries, delineating 135 CIDs-like regions ranging in size from 5 to 125 kb (magenta signal, **Figure 1e,f**; Supplementary data Table 1). Among those, 22 overlap with those called with the DI analysis of the 5kb binned contact map (blue signal, **Figure 1e,f**) and enriched HEG annotations. The remaining 113 positions correspond to less expressed genes (**Figure 1f**). Altogether, these results suggest that the chromosome, rather than being structured into large self-interacting regions, is organized by a succession of TIDs, reminiscent of those observed in budding yeast (Hsieh et al. 2016). These small transcribed domains are separated by non-transcribed regions depleted in local Hi-C contacts in the maps.

### Expression of a single TU is sufficient to imprint a domain on Hi-C contact maps

To further understand the nature of the transcription-dependent, short-range contacts increase observed in the high-resolution contact maps, we designed an artificial inducible system. A T7 promoter was inserted at the LacZ locus, facing towards the *ter*. The T7 RNA polymerase is specific to its own promoters, and was put under the control of the inducible arabinose promoter (**Figure 2**). Upon arabinose addition, a strong, dense signal appears on the Hi-C map, originating at the pT7 position and propagating towards the Ter over ~ 70 kb (**Figure 2a,b**). Chromatin immunoprecipitation of the T7 RNA polymerase showed a strong enrichment at the pT7 (Supplementary Data Figure 2a), whereas RNA seq analysis further confirmed the strong induction of this artificial TU (**Figure 2b**). Since the T7 RNA polymerase is insensitive to the bacteria RNA polymerase inhibitor rifampicin, we reasoned that treating the cells with the drug should result in the unveiling of a single transcriptional unit (Tabor and Richardson 1985). Indeed, the normalized contact map of exponentially growing cells treated with rifampicin display a clear, discrete signal at the level of the T7 promoter, further magnified when plotting the ratio between the maps of cells treated with rifampicin but with or without T7 induction (**Figure 2a**, bottom). In absence of neighboring transcription, the T7 promoter resulted in a longer transcription track covering ~ 110kb (**Figure 2a,b**). The difference between the transcription track length in cells treated or not with rifampicin suggests that T7 transcription is limited by neighboring transcription. Consequently, this system allows to magnify a signal emanating from a single TU. Magnification of the induced T7 region from the normalized wt +rif map reveals two types of contact patterns at the induced promoter: a “stripe”, extending from the TSS, and a thick, short range contact pattern extending across the transcription and T7 RNApol deposition tracks, as determined by RNA-seq and ChIP-seq, respectively (**Figure 2b**; Supplementary Data Figure 2a). Both signals were observed upon inversion of the gene (Supplementary Data Figure 2a). The thick pattern, but not the stripe, is strongly reminiscent of that observed from the pile up plots of highly expressed TSS of the native genome (**Figure 1c**). The transcribed region is covered with polysomes, and thus most likely translated (Supplementary Data Figure 3). However, the contact signal appeared independent of translation, as two stop codons introduced downstream the promoter pT7*lacZ^2Xstop^* did not suppress it (Supplementary Data Figure 3).

**Figure 2:**
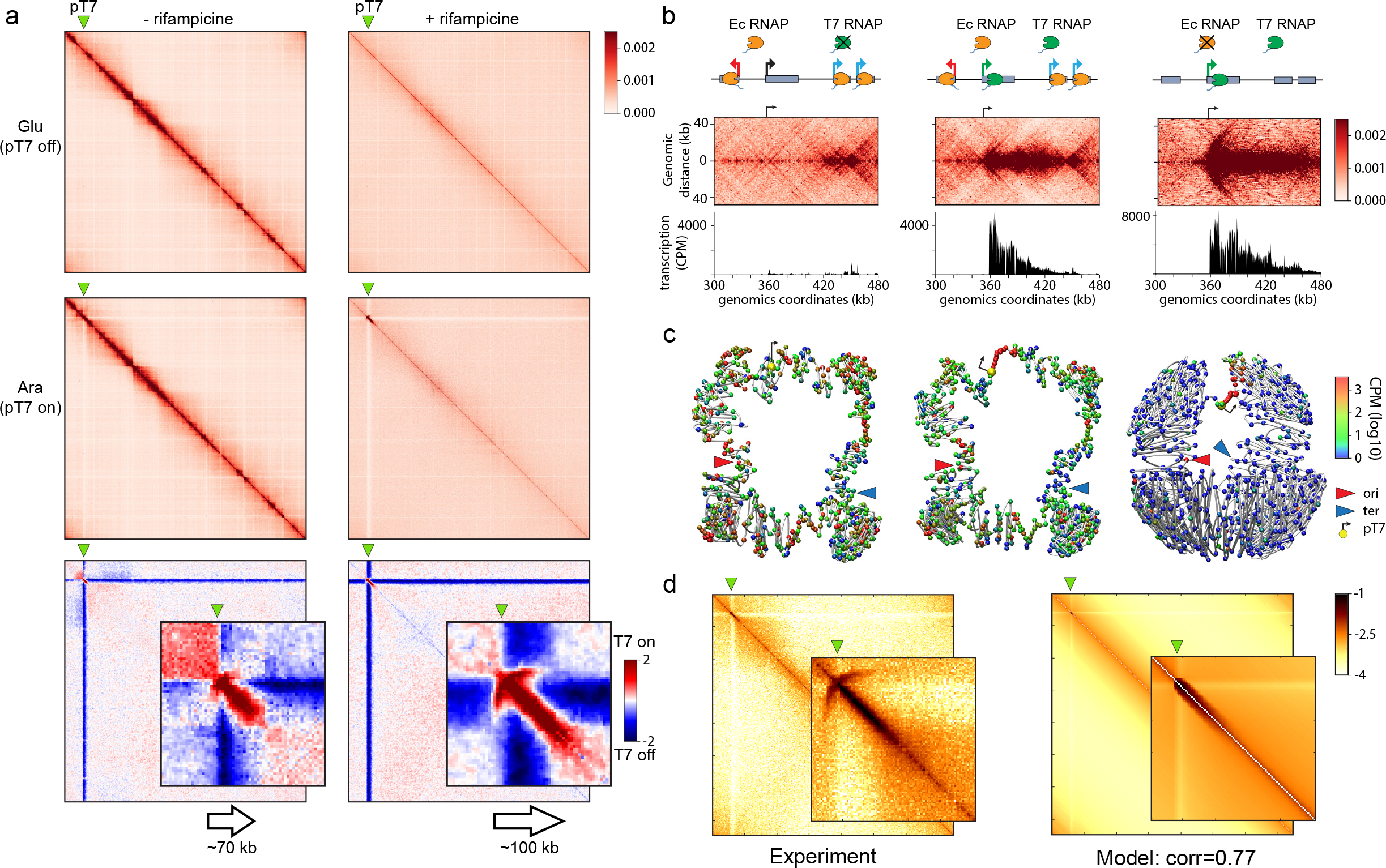
Contact profile of single, active transcription unit within an entire genome. **a,** Hi-C contact maps (bin: 1kb) of *E. coli* chromosome carrying a single T7 promoter (green triangle), in absence (left) or presence (right) of rifampicin. Top panels: cells grown in glucose media, when the T7 RNA polymerase is not expressed. Middle: cells grown in presence of arabinose, with expression of the T7 RNA polymerase. Bottom: log2 ratio contact maps with and without induction of the T7 promoter. Magnification of the T7 promoter region, represented using Serpentine flexible binning (Methods)(Baudry et al. 2020). **b,** Magnification of the T7 promoter in the normalized contact maps, with and without induction and in presence and absence of rifampicin. From left to right: T7 promoter off, no rif; T7 on, no rif; T7 on, + rif. The corresponding RNA-seq tracks (CPM) are plotted under the maps. **c,** 3D representation using Shrek of the corresponding 2D contact maps of the *E. coli* bacterial chromosome in the different conditions. The green, red and blue arrows represent the pT7, ori and ter positions respectively. **d**, Modelisation of the Hi-C contact maps using the RNA polymerase distribution on the genome and using the second model (Methods).

### Modeling transcription induced domains

The 3D representation of the contact map using the shortest-path reconstruction algorithm ShRec3D further shows how the highly transcribed T7 TU contact map translates into a discrete structure within the chromosome that insulates flanking regions (**Figure 2c**) (Lesne et al. 2014). While this insulation effect can be seen both in the case for which transcription is active and inactive it appears stronger when rifampicin is present since the overall structure of the chromosome is more intertwined, due to a strong increase of long-range contacts over short-range contacts.

To gain more quantitative insight into the link between transcription and increased short range contacts, we developed a probabilistic modeling approach to emulate the observed contact map under two different assumptions. In the first hypothesis, the increase in short-range contacts is due to the existence of preferential contacts between T7 RNA polymerases. In the second model, we added an insulation effect for each polymerase such that the contact probability between two polymerases decreases if another polymerase is present between them. The models take as inputs the experimentally measured decay in contact frequency with increasing genomic distance and the ChIP deposition profile of T7 RNApol. The only parameter is the maximum T7 RNApol occupancy, set to get the highest correlation between the experimental Hi-C and the model contact maps. The best result (correlation of 0.77) was obtained for the second model and a parameter of value of 0.15 (Supplementary Data Figure 2c; compared to a maximum correlation of 0.67 for model 1 in Supplementary Data Figure 2d). The extended, thick pattern correlates nicely with the experimental T7 RNApol occupancy, suggesting that crosslinking of trains of consecutive RNA polymerase along the transcribed track are responsible for generating the contact pattern observed (**Figure 2d**). These results suggest that the thick motif observed in TIDs of Hi-C contact maps corresponds to trains of RNA polymerases that each have a cumulative local insulating effect.

### Interactions between adjacent T7 RNApol induced domains

We next combined pairs of transcribed T7 units (pT7lacZ and pT7mCherry) to further characterize the potential structural interplay between two neighboring genes. The second pT7 was introduced at either 60 or 100 kb upstream of pT7lacZ, either in collinear, convergent, or divergent orientation (**Figure 3**, panel i). Exponentially growing cells were induced for T7 RNA pol using arabinose, treated with rifampicin, and processed with Hi-C and RNA-seq. In all cases, we observed an excellent correlation between the short-range contacts, transcription tracts, and T7 RNA pol as quantified using ChIP-seq (Spearman correlation between 0.62 and 0.91) (**Figure 3**, panels i, ii, iii). However, the stripe pattern appears affected by the orientation of the promoters with respect to each other. Upon induction, the two promoters positioned in divergent orientations and separated by 100 kb displayed similar contact patterns, i.e. a stripe and the globular short-range tract (**Figure 3b**, panels i and ii). The stripe pattern vanished when the distance separating the promoters was shortened (60 kb), resulting in a self-interacting domain of enriched contact positioned in-between the two genes (**Figure 3c**, panels i and ii). In contrast, the two genes in convergent orientation resulted in the two transcription tracks abruptly ending at mid-distance, resulting in a sharp boundary right in-between the two promoters (**Figure. 3d**, panels i and ii). When positioned in colinear orientation, the clearly visible stripe of the pT7lacZ promoter appeared strongly reduced if not entirely suppressed by the incoming transcription tract of the upstream pT7mCherry (**Figure 3e, f,** panels i and ii).

**Figure 3:**
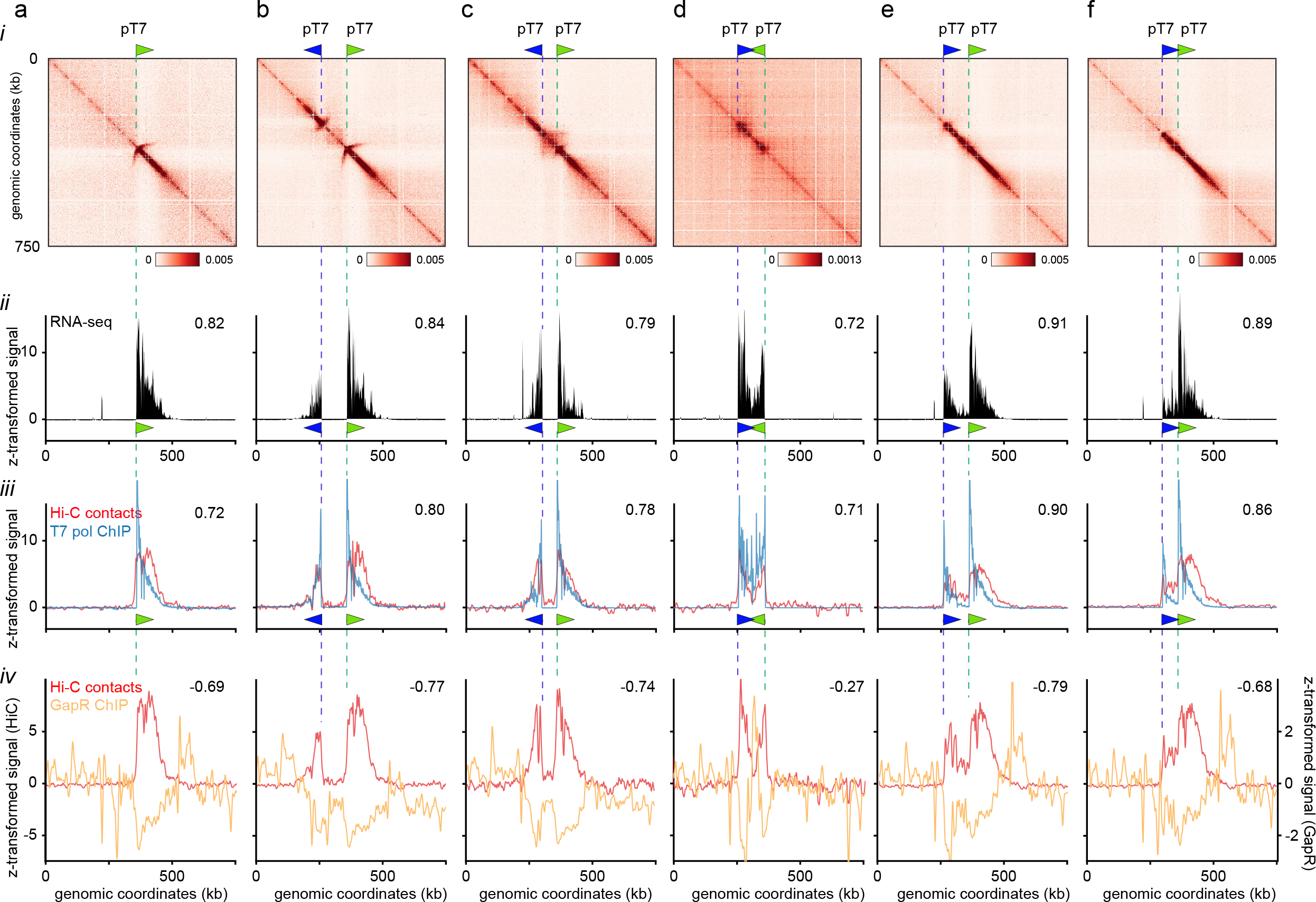
Supercoiling and contacts resulting from combination of pairs of transcription units. Genomic characterization of chromosomal regions carrying pairs of pT7 promoters in different orientations. From top to the bottom: Hi-C contact map (i; bin = 1kb), RNA-seq track (ii; in CPM), T7 RNA polymerase ChIP-seq track (iii, blue curve) and short-range Hi-C contacts (iii, red curve), and GapR ChIP-seq revealing positive supercoiling (iv, yellow curve) and short-range Hi-C contacts (iv, red curve). Values on the top right corner of each panel are the Spearman correlation coefficients of the track with the short-range Hi-C contacts. All tracks are z-transformed.

### Characterization of the stripe feature of the T7 RNApol induced domains

Transcription is known to promote the formation of positive and negative supercoils in front of and behind the RNA polymerase (Liu and Wang 1987; Visser et al. 2022). An attractive interpretation of the T7 TU domains plasticity with their genomic organization is that in collinear orientation, the positive supercoils of the first transcribed track will offset the negative supercoils generated by the second one. Since the stripe is abrogated only in this orientation, while transcription and occupancy of the T7 pol DNA are conserved, it is probable that this structure corresponds to negative supercoiled signal generated at the TSS. This hypothesis is supported by the fact that novobiocin-induced inhibition of DNA gyrase, which actively introduces negative supercoils into DNA (Levine et al. 1998), resulted in a strong disruption of genome-wide Hi-C contacts, including loss of the stripe at the level of the T7 unit (Supplementary Data Figure 2b). To measure the supercoiled nature of the DNA template in this region, we quantified the deposition of GapR, a *Caulobacter crescentus* protein recently introduced as a marker of positive supercoiling along bacterial and yeast chromosomes (Guo et al. 2021). As expected, a small enrichment of GapR was observed at the single T7 RNA track end (**Figure 3a**, panel iv). A depletion was observed in-between genes in divergent orientations (**Figure 3b, c**, panel iv), whereas GapR was enriched in-between the two genes positioned in convergent orientations, (**Figure 3d**, panel iv). In colinear orientation, no enrichment was seen after the first gene, in agreement with the suppression of the positive supercoils by the neighboring negative one (**Figure 3e, f**, panel iv). However, a strong enrichment was observed after the second gene. We therefore propose that the stripe pattern is the Hi-C signature of a negative supercoiled structure positioned in the 5’ region of the gene. These observations are in agreement with simulated and experimental data pointing at a preferred positioning of RNA polymerase at the apical positions of supercoiling loops (ten Heggeler-Bordier et al. 1992; Racko et al. 2018). A fine observation of the signal suggests that, indeed, the T7 promoter is positioned in the middle of the stripe. On endogenous genes, this pattern would be either two small to be visualized, erased by neighboring supercoiling, or both.

### T7 RNApol induced domains impose mechanical constraints on adjacent chromatin

The T7 system provides a distinct opportunity to investigate the impact of a single TU on the mobility of its neighboring regions. To monitor the influence of T7 transcription on the mobility of chromosome loci, we used strains carrying fluorescently labeled lacO20 arrays inserted at four positions either upstream or downstream of the T7 promoter (**Figure 4a**; Methods). We compared individual foci dynamics with or without T7 induction, in the absence or presence of rifampicin, by recording their position every second for 120□s (**Figure 4b**; Supplementary Data Figure 4). For each trajectory, we plotted the diffusion coefficient (D_c_) as a function of the slope (α) of the MSD versus time interval. D_c_ accounts for local viscosity that may vary along the genome while α is indicative of the nature of the locus movement. α□=□1 describes normal diffusion, whereas α□<□1 is sub-diffusive (constrained) and α >□1 points at super-diffusive (directed) movement (Methods). On average, we observed a linear anti-correlation between α and D_c_, with D_c_ decreasing while α increases (linear regression log (D_c_) ≈ −A log (α), with A depending on the locus and conditions). For α lower than 0.35, we observed a second population of trajectories with high D_c_ (> 2×10^−3^ μm^2^.sec^−1^, i.e. log (D_c_) ≈ −0.22; top left corner delimited by the dotted lines). For the loci close to the T7 TU (*betT*, *tauB* and *ecpR*), T7 activation correlated with a higher proportion of foci with a fast diffusion and low α (**Figure 4b**, pink curves; Supplementary Fig 4). In contrast, a focus positioned 2 Mb away at the *yqeK* locus did not show significant changes (**Figure 4b**). In the presence of rifampicin, the *yqeK* control locus displayed a narrower range of D_c_ and α, suggesting that transcription contributes directly to chromatin dynamics. By contrast, mobility of the T7 proximal loci was unchanged by rifampicin (**Figure 4b**; Supplementary Figure 4c), suggesting that transcription directly influences chromatin dynamics. In presence of rifampicin, the influence of T7 induction was manifest for the *ecpR* locus. However, for loci close to the T7 TSS (*betT* and *tauB)* T7 induction in the presence of rifampicin had limited impact on DNA mobility suggesting that transcription leakage in repressive conditions might be sufficient to stimulate chromatin mobility. These observations confirm that transcription has a direct but local influence on chromatin mobility: it simultaneously increases compaction of the region, in agreement with the Hi-C data, while enhancing its mobility.

**Figure 4:**
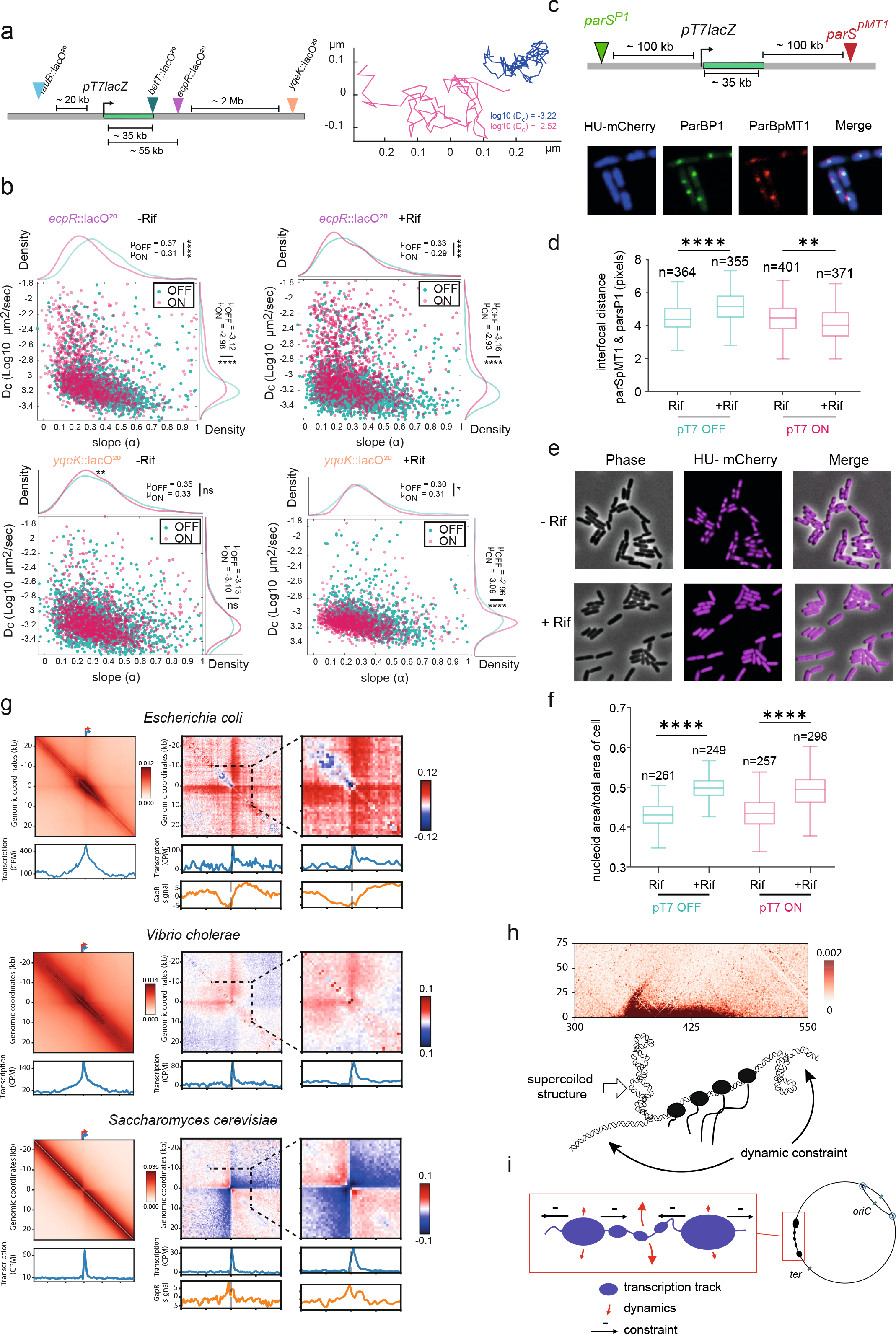
Dynamic influence of the T7 transcription unit. **a,** Positions of the four lacO arrays inserted in the vicinity of the T7 promoter (left). Two trajectories of the same locus representative of the two populations of diffusion coefficients (D parameter) (right). **b,** Scatter plots of D_c_ as a function of the MSD α for ~1,000 trajectories of fluorescently labeled loci upon induction of the T7 TU in the absence or presence of rifampicin. Top panels: *ecpR* locus positioned 50kb from T7 TU TSS. Bottom panels: *yqeK* locus positioned 2Mb from T7 TU, the mean (μ) of each distribution is indicated. Statistical differences are measured by an Anova Kruskal-Wallis test with Bonferroni correction, * <0.033, ** < 0.0021, **** <0.0002, ****<0.0001. The distribution are well fitted with Log_10_ (D_c_)= A* Log_10_(α) −B, A is indicated below the scatter plots, B ~-3. **c,** Positions of two fluorescently labeled loci (*par*^P1^ and *parS*^pMT1^) 100 kb upstream and downstream of the T7 TU and representative image. **d,** Distribution of interfocal distances computed for each condition over >350 cells. Data were analyzed using a Student test **e-f,** Representative images of HU-mCherry-labeled nucleoids under the different conditions, and corresponding distributions of % nucleoid occupancy in the cell. **g,** *E. coli*, *V. cholerae*, and *S. cerevisiae* genes pileups. Left: pileup for each species of 50 kb windows contact maps centered on TSS of the 10% most transcribed genes. Below: corresponding RNA-seq pileup profiles. Right panel: log ratio of pileups of 50 kb windows contact maps centered on the 20% most transcribed TUs centered on their TSS, over pileup of random 50 kb windows. 20 kb window magnification is shown. The GapR ChIP-seq track is shown for *E. coli* and *S. cerevisiae* (Guo et al. 2021). **h,** Schematic interpretation of the contact patterns observed in the single TU contact map. **i,** Schematic representation of the proposed nucleoid structuring into a mosaic of small, locally constrained 3D transcriptional units.

Next, we tested with DAPI and pairs of fluorescent *parS*/ParB tags positioned 100 kb upstream and downstream the T7 promoter whether transcription influences nucleoid folding on a larger scale (**Figure 4c-f**). In the absence of rifampicin, the interfocal distance between the two foci remains the same with or without T7 activation (**Figure 4d**). When the T7 TU was silent, the presence of rifampicin increased significantly the distances between the *parS*^P1^ and *parS*^pMT1^ foci, in agreement with the accompanying general nucleoid decompaction (**Figure 4e, f**). However, when the T7 TU was active the distance between foci was not affected by rifampicin (**Figure 4d**). This result shows that transcription locally enhances the compaction of neighboring regions, maintaining the two foci closer together, presumably through the formation of supercoiled loops.

These experiments reveal that in *E. coli* DNA mobility is intricately linked to local transcription, which imposes a mechanical constraint on neighboring loci by moving them closer to each other (Germier et al. 2017; Gu et al. 2018).

## Discussion

The thinner grain scale made available by resolution improvements combined with the single transcription unit analysis suggests that transcription in bacteria imposes multilevel constraints. We demonstrate that the large CIDs identified from the long HEG are the tip of a much more general phenomena, also visible in the high-resolution Hi-C contact maps of another bacterium (*Vibrio cholerae*, **Figure 4g**). The picture emerging from the present analysis is that transcription shapes bacterial chromosomes locally. Transcription locally stimulates the formation of globular domains (TIDs), interaction between adjacent genes or operons and the emergence of a negatively supercoiled stripe/loop at the 5’ end of TUs (**Figure 4h-i**). These features are influenced by the transcription level and the genomic context, suggesting that the chromosome is not under a homogeneous amount of transcription-mediated mechanical constraints. The relationship between TIDs and long-range chromosome organization (e.g. plectenomic loops(Deng et al. 2004; Postow et al. 2004), macrodomains (Valens et al. 2004), supercoiling domains (Visser et al. 2022)) is not yet known. For instance, HEG are frequent in the *oriC* proximal part of the genome, and the resulting TIDs may influence the long range *oriC* interaction revealed by Hi-C(Lioy et al. 2018; Marbouty et al. 2015), recombination (Valens et al. 2004) and imaging (Marbouty et al. 2015) or supercoiling of this region (Visser et al. 2022). Finally, although Hi-C is not suitable to measure contacts between repeated sequences, a careful examination of the high-resolution map points at weak contacts between regions flanking ribosomal RNA operons, confirming earlier imaging observations 23. This pattern falls within the more general propensity of all adjacent expressed sequences to contact each other more frequently, and thus this behavior would not be specific but only magnified at ribosomal DNA operons.

Sub-kb resolution Hi-C reveals plaid-like patterns with interactions between neighbor TUs (Figure 1), a new feature of bacterial chromosome folding. Since these distant contacts (≈ 20 – 40 kb) can involve protein coding genes (membrane and cytoplasmic proteins) and tRNA regulated by different transcription factors and different sigma factors (**Figure 1c**), they may only rely on transcription. Several hypotheses may explain this phenomenon. First, the increased mobility of transcribed units (**Figure 4a–d**), in association with their relocalization to the nucleoid periphery (Stracy et al. 2015), could explain inter-transcriptional unit contacts. Second, the proximity of transcribing RNA polymerases may favor protein-protein interactions as biomolecular condensates (Ladouceur et al. 2020). Third, RNA production may locally reduce solvent quality of the cytoplasm and drive local chromosome deformation as proposed by the group of Christine Jacobs-Wagner (Xiang et al. 2021). Alternatively, one cannot exclude that contacts between adjacent transcribed regions are also regulated by condensin loop extrusion, an active process that expands DNA loops (Brandão et al. 2019). Loops would extend until they encounter actively transcribed regions that would act as roadblocks or extrusion slowing zone, resulting in enriched contacts between them. Future experiments will be required to evaluate the contribution of these elements for the folding of transcribed units.

It is tempting to propose that the globular signal overlapping TU, and the inter TU contacts, relies on a single biophysical property of transcribed chromatin. However, the 5’ stripe of T7 TU that is sensitive to genes organization is most likely linked to transient supercoiling waves. It is manifest at the T7 TSS but rarely on the much smaller endogenous genes, and reduced in normal compared to rifampicin conditions, suggesting the strong activity of the T7 RNApol exacerbate an otherwise more discrete signal. Therefore, it probably reflects a dramatic underwounding of the DNA following T7 RNApol transcription that Gyrase fails to counteract. Similarly, the signature of the 25 kb twin-supercoiled domains recently described using psoralene crosslinking is only detectable around highly expressed genes (i.e. ribosomal operons (Visser et al. 2022). At homeostasis, we propose that the constraints imposed by transcription along the fiber will balance each other allowing compaction, organization and dynamics of the chromosome. However, transiently these constraints could have multiple consequences for DNA transactions, including transcription, DNA repair, and segregation, as well as contribute to the regulation of the extrusion of large DNA loops by bacteria condensins as they travel along the chromosome (Mäkelä and Sherratt 2020; Wang et al. 2017; Brandão et al. 2019).

In Eukaryotes, transcription shapes chromosome architecture but the contact patterns differ, with active genes delineating clear boundaries in contact maps for instance in *S. cerevisiae* (**Figure 4g**), as shown by past and recent work (Hsieh et al. 2016; Banigan et al. 2022). This pattern appears modulated by structural maintenance of chromosome complex (SMC) DNA translocase activity (Racko et al. 2018; Banigan et al. 2022). The presence of nucleosomes in eukaryotes and in some archaea is also expected to thicken the contact pattern at short distance, therefore blurring the crispier signal observed in bacteria. Nevertheless, the underlying constraints unveiled in this work imposed by transcription on the DNA sequence stand to be a fundamental aspect of chromosome biology.

## Acknowledgements

We thank Michael T. Laub and Monica Guo for sharing with us the FLAG-tagged GapR construct. This research was supported by the European Research Council under the Horizon 2020 Program (ERC grant agreement 771813) to RK, by Agence Nationale de la Recherche (ANR Hiresbac) to RK, JM and OE, and by the QLife program to RK and OE. We thank all our colleagues from the laboratory régulation spatiale des génomes for fruitful discussions. We especially thank Axel Cournac and Marianne de Paepe for early discussion on the project, and are grateful to Aurèle Piazza and Frédéric Beckouët for comments on the manuscript.

## Author contributions

Conceptualization: CC, OE and RK. Methodology: CC, AB, JM, OE, RK. Investigation: CC, with contributions from EA. Formal analysis: AB, JM, CC, OE. Data Curation: AB, RK. Visualization: AB, RK. Writing - original draft, RK. Writing - Other drafts & Editing, RK, OE, AB. Supervision: OE, RK. Funding acquisition: OE, JM, RK.

## Declaration of interest

The authors declare no competing interests.

## Data Availability

Sample description and raw sequences are accessible on the SRA database through the following accession number: XXX.

## METHODS

### Media culture conditions and strains

All strains used in this study are derived from the BW25113 E. coli strain and are listed in the Supplementary Table 2. All strains were grown in minimal media A (0.26 M KH2PO4, 0.06 M K2HPO4, 0.01 M tri sodium citrate, 2mM MgSO4, 0.04 M (NH4)2SO4) supplemented with 0.2% of casamino acids and 0.5% of glucose at 37°C. The strains were grown with 0.2% arabinose for 1h to induce T7 RNA polymerase expression under the control of the PBAD promoter.

### Drugs and antibiotics

Rifampicin was used for 10 min at a 100 μg/ml working concentration to inhibit transcription.

### Hi-C procedure and sequencing

Cell fixation with 3% formaldehyde (Sigma-Aldrich, Cat. F8775) was performed as described in Cockram et al. (2021) (Cockram et al. 2021b). Quenching of formaldehyde with 300 mM glycine was performed at 4°C for 20 min. Hi-C experiments were performed as described in Cockram et al. (2021). Samples were sonicated using Covaris (DNA 300bp).

### ChIP- and RNA-seq experiments

Chromatin immunoprecipitation was performed as described (Cockram et al. 2015). Briefly, overnight cultures were diluted to OD_600nm_= 0.01, grown until OD_600nm_= ~0.2 – 0.25, diluted and crosslinked using formaldehyde (Sigma-Aldrich; final concentration of 1%) for 10 minutes at 22.5°C. Formaldehyde was then quenched by adding 2.5M glycine (final concentration: 0.5 M), for 10 minutes at room temperature (e.g. 19 – 22°C). Cells were collected by centrifugation at 1,500 × g for 10 minutes and washed three times in ice-cold 1X PBS. The pellets can be stored at −80 °C or used straight away. A pellet was then resuspended into 500 μl of 1X TE buffer, supplemented with 5 μl of ready-lyse lysozyme, and incubated with shaking at 37 °C for 30 minutes. 500 μl of 2X ChIP buffer (50 mM HEPES-KOH pH 7.5, 150 mM NaCl, 1 mM EDTA, 1% Triton X-100, 0.1% sodium deoxycholate (DOC), 0.1% SDS 1X Roche Complete EDTA-free protease inhibitor cocktail) was then added and the sample transferred to ice. The sample was transferred to a pre-chilled 1 mL Covaris tube (Covaris), and sonicated using Covaris S220 for 7 min (settings as followed: target size, 200 – 700; PIP 140; DF 5%; CPB 200). 100 μl of the sample was removed as input and stored at 20. Immunoprecipitation was performed overnight under rotation at 4 using 1/100 T7RNA antibody (Biolabs CB MAB-0296MC) and antiflag (Sigma F1804 and F3165). Immunoprecipitated samples were incubated with Protein G Dynabeads (Invitrogen) with rotation for 2 min at room temperature. The tube was washed three times with 1X PBS with 0.02% Tween-20 using the Dynamag magnet setup. The beads were resuspended in 200 μl TE buffer with 1% SDS and 1 μl RNAseA (10 mg/ml) and 1 μl proteinase K (20 mg/ml). Samples were incubated at 65 for 10 h to reverse the formaldehyde crosslinking. The beads were removed using the Dynamag magnet and DNA of the supernatant purified using Qiagen Minelute PCR purification kit using two elution steps. DNA was eluted into a 50 μl TE buffer and stored at −20 until further processing.

### RNA-seq

Total RNA was extracted from E. coli using the Nucleospin RNA Extraction Kit (Macherey-Nagel) according to manufacturers’ instructions. DNAse was depleted using an additional DNase treatment with Turbo DNase (Thermo Fisher). The DNAse was inactivated and RNA purified by a phenol-chloroform extraction (pH 4.5, Amresco) and ethanol precipitation. The RNA was then resuspended in DEPC-treated water. Ribosomal RNA depletion was done using Ribo-Zero magnetic beads according to manufacturer protocol (Illumina). cDNA library preparation was performed following standard protocols. Briefly, RNA was fragmented using the NEBnext mRNA first and second strand synthesis kits (NEB). One to three biological replicates were generated for each condition and on average ~10 million reads were generated per sample.

### DNA libraries preparation

For Hi-C, RNA-seq and ChIP-seq libraries, preparation of the samples for paired-end sequencing was performed using Invitrogen TM Colibri TM PS DNA Library Prep Kit for Illumina according to manufacturer instructions. The detailed protocol is available in Cockram et al. (2021) (Cockram et al. 2021b). All libraries used or generated during the course of this study are listed in Supplementary Table 3.

### Gradient preparation of *E. coli* polysomes

To preserve the polysomes, cultures of E. coli are incubated with 100μg/mL of chloramphenicol before centrifugation. Fresh cell paste (0,7g) was homogenized in the buffer (150mM NH_4_Cl, 10mM MgCl_2_, 2mM Tris-Cl pH 7.5, 10μM PMSF, 0.2μg/mL chloramphenicol, Complete EDTA free, RNAsine) at a 1:2 (w:v) ratio and set aside for 20 min at 4°C. Disrupt cells using FastPrep® sample preparation system and lysing matrix B tubes (2mL) containing 0.1mm silica beads. Add sodium deoxycholate (1% final), DNase I to a final concentration of 2μg/mL (20U/ml) and let 30mn on ice, then clear the lysate of cell debris by centrifugation at 4°C using top bench centrifuge for 20mn and a second centrifuge of 5mn. Divide the supernatant equally, and treat one part by adding EDTA (70mM) and incubate on ice for 30mn. Layer the fractions (600μL) on top of 10mL sucrose gradient (10-40%) and centrifuge for 2.5hr at 4°C in SW41Ti rotor at 35,000rpm (151,000g). Gradients are next fractionated by collecting 500μL fractions. To analyze RNA, 170μL of each fractions is mixed to 400μL of RNAse-free water and 570μL of phenol, vortexed and centrifuged to extract RNA from proteins, then aqueous supernatant is precipitated with CH3COONa, Glycogen and isopropanol. Collected RNA present in each fraction is next analyzed in agarose gel.

### Processing of reads and Hi-C data analysis

Reads were aligned with bowtie2 v2.4.4 and Hi-C contact maps were generated using hicstuff v3.0.3 (Matthey-Doret et al. 2020) (https://github.com/koszullab/hicstuff) with default parameters and using HpaII enzyme to digest. Contacts were filtered as described in Cournac *et al.* (2012) (Cournac et al. 2012), and PCR duplicates (defined as paired reads mapping at exactly the same position) were discarded. Matrices were binned at 0.5, 1, 2 or 5kb. Balanced normalizations were performed using ICE algorithm (Imakaev et al. 2012). For all comparative analyses, matrices were down-sampled to the same number of contacts. Comparison between matrices were done using log2 ratio and serpentine v0.1.3 (Baudry et al. 2020) for flexible binning. Serpentine was used with 5kb binned matrices, with 25 iterations and a threshold of

100. The Hi-C signal was computed as the contacts between adjacent 5kb bins as described in Lioy *et al*. 2018 (Lioy et al. 2018). In order to compare this signal with other genomics tracks, we binned it at the desired resolution and z-transformed it.

### Borders detection

To detect the borders we first used the directional method as described in Lioy *et al.* 2018 (Lioy et al. 2018). The directional index is a statistical parameter that quantifies the degree of upstream or downstream contact bias for a genomic region (Dixon et al. 2012). For each bin, we extracted the vector of contacts from the correlation matrix between that bin and bins up to a window size in both left and right directions. To assess if the strength of interactions is stronger with one direction relative to the other we used a paired t-test between the two vectors. A p-value of 0.05 was used as a threshold to assess a statistical significant difference. The directional preferences for the bin along the chromosome are represented as a bar plot with positive and negative t-values shown as red and green bars, respectively. We trimmed the bars of the bins with t values below −2 or above 2 (corresponding to a p-value = 0.05). At the borders identified in the contact matrices, the directional index changes from negative to positive t-values. The implementation of the code is available at https://github.com/ABignaud/bacchus and it’s based on the one used for Lioy et al., 2018 (Lioy et al. 2018). The DI method is depending on the binning resolution and on the window size. At small window size, it misses the larger domains visible at larger scale and at large window size it only finds the larger domains. Moreover, the resolution impacts on the performance of the DI, at low resolution it cannot find the smallest domains which are merged in few bins and at high resolutions it starts to be noisy as the resolution directly impacts the width of the vectors used to compute the DI. In our study, we decided to use an insulation score method to improve the borders detection at higher resolution. For our analysis, we developed a python implementation (https://github.com/koszullab/bacchus) of the HiCDB algorithm (Chen et al. 2018). This method allows multiple window sizes, which reduces the dependence between the window size and the size of the detected domains. Furthermore, it does not depend on the resolution of the matrix, which allows for efficient detection of boundaries even at high resolution. We used the 1kb resolution contact map with 10, 15, 20, 25 and 30kb windows (Supplementary data table 1).

### Pileup analysis

The Hi-C contacts were built and normalized as explained before at a resolution of 500bp. For each gene we extract a 100kb matrix centered on the start codon of the gene. For reverse genes, we flip the matrix to have the centered genes pointing always in the same direction. The pileup plot is the average of all the extracted windows, without taking into account the white lines (i.e. bins with less than the median minus three times the median absolute deviation are considered as white lines). To select active genes, we select a fraction of the most transcribed genes (values in Reads Per Kilobase per Million (RPKM)) as the active genes. For the transcription units analysis, to center our windows on the first transcribed genes, we selected active genes only if there are no other active genes in the 3kb upstream of the start codon of the gene. To compare the pileups of the first transcribed genes with the non-coding or non-transcribed regions, we calculated the ratio between the pileup of the first transcribed genes and the pileup of random windows taken from the same region (center on a random position within 100kb around the gene). We chose to use random regions instead of the pileup of non-coding genes or the expected matrix (matrix corresponding to the contacts of the genomic distance law) to avoid having a bias of the region where we extract the active genes.

### Detection of contact bundles (i.e. TIDs) along the main diagonal

To detect contact bundles on the main diagonal, we used a convolution kernel on the balanced matrix. The method is implemented in (https://github.com/koszullab/bacchus). We used a computer vision approach similar to the program Chromosight ^45^, we use a convolution kernel, describing a given pattern, as a template to detect the local similarity with it. Here, we aim at detecting the bundles on the main diagonal of the matrix. To detect them, we build a gaussian kernel of size n as follow (n=5 in our study):

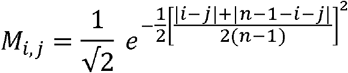

By computing the convolution product between each local image centered on each bin of the main diagonal and the kernel, we obtain a convolution score. The higher the score is, the closest the local image is to the kernel and the more likely it is to be a bundle. To remove the effect of local regions, we remove the second envelope of the signal as it’s described in the HiCDB ^24^ insulation score algorithm. Finally, the borders of the bundles are detected by taking each peak of the local convolution score superior to the median of the local convolution score. The bundle region is then extended until the value gets inferior to one third of the peak.

### RNA-seq processing

Processing is done using tinymapper v0.9.14 (https://github.com/js2264/tinyMapper) with default RNA parameters. The reads are mapped with bowtie2 v2.4.4, PCR duplicates are filtered using samtools v1.14 and Count Per Million (CPM) is made with bamCoverage v3.5.1. We used only the unstranded signal, and binned it depending on the displaying resolution. For the comparisons with other signals, a z-transformation is done.

### ChIP-seq processing

Processing of the ChIP-seq of T7 RNA polymerase and GapR is done using tinymapper v0.9.14 (https://github.com/js2264/tinyMapper) with default ChIP-seq parameters without input. The reads are mapped with bowtie2 v2.4.4, PCR duplicates are filtered using samtools v1.14 and CPM is made with bamCoverage v3.5.1. For the GapR-seq, we do a gaussian blur of the signal with the gaussian_filter1d function from scipy v1.7.3 with ‘wrap’ mode and sigma value of 2500, as described in Guo *et al*. 2021. The data is then binned at the displaying resolution and z-transformed to compare it to other signals.

### Imaging and analysis

Cells were grown similarly to Hi-C samples (above). One hour after arabinose induction of T7 RNA polymerase, 2 mL of cells were pelleted and resuspended in 50 μl of fresh medium. Three drops of 2μl are deposited on a freshly made agarose pad (1× supplemented medium A, 1% agarose) incubated 30 min in the microscope incubation chamber at 37°C and imaged for 120 sec every sec for 100msec. For foci mobility analysis, imaging was performed on a Nikon Eclipse Ti inverted microscope equipped with a Spinning-Disk CSU-X1 (Yokogawa), an EM-CCD Evolve 512*512, magnification lens 1.2, pixel size: 13.3 μm * 13.3 μm camera at 600-fold magnification. Focal plane was maintained during acquisition using Nikon Hardware autocus. Illumination and acquisition was controlled by Metamorph. Time series images were registered using Stackreg (http://bigwww.epfl.ch/thevenaz/stackreg/) and analyzed with the MOSAIC suite (https://git.mpi-cbg.de/mosaic/software/bio-imaging/MosaicSuite) as FIJI plugins. MSD (a) and Dc distribution were analyzed and plotted with MATLAB. An average of 1,000 trajectories were analyzed for each replicate. Experiments were performed twice for each strain and conditions. For interfocal distances and nucleoid compaction measurements, cells were observed live on agarose pad on a thermo-controlled stage with a Spinning disk (Yokogawa) W1 system mounted on a Zeiss inverted confocal microscope and a C-MOS Hamamatsu 2048*2048 / pixel size : 6.45*6.45 μm camera at 630-fold magnification. The position of foci in the cell in each condition was analyzed with the ObjectJ plugins of ImageJ https://sils.fnwi.uva.nl/bcb/objectj/. Two color localization was performed with 7 couples of tags forms with combination of *crl* parS P1 and *cynX* parS pMT1 tags (Vickridge et al. 2017). An average of 600 cells were analyzed per strain and condition.

### Modeling approach

We devised a simple model to reproduce the contact maps obtained experimentally under a few hypotheses. We start our approach by computing the contact probability decline with increasing genomic distance from experimental data p(s). We then make the hypothesis that two different types of contacts are found in the experiments: contacts mediated by polymerases and contacts mediated by other proteins. We assume that the proportion of contacts mediated by polymerases at bin i is C_i_, where C_i_ is the normalized experimental Pol-ChiP signal. To normalize the signal, we define its maximum value as ε, which is between 0 and 1. The proportion of contacts mediated by other proteins at bin i is then simply 1-C_i_. We then compute the contact probability between any couple of bins i and j using two different models:

- In model 1, there is a preferential interaction between polymerases so that the contact frequency is proportional to: p(s_i,j_) × (C_i_C_j_ + (1 − C_i_)(1 − C_j_)
- In model 2, there is a preferential interaction only between consecutive polymerases. The idea behind this model is that polymerases also act as contact insulators. The contact frequency is then modified from model 1: p(s_i,j_) × (m × C_i_C_j_ + (1 − C_i_)(1 − C_j_) with 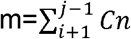representing the insulation factor, which is the amount of polymerase that is found between bins i and j.

After all contact probabilities have been computed for each model, the contact matrix is normalized so that the sum of each line and each column is equal to one so that it corresponds to contact probabilities. The Spearman rank correlation is then computed between the experimental map and the model map is then computed to find the best value for epsilon and to compare the relevance of each of the two models.

### Data and software accessibility

The accession number for the sequencing reads reported in this study is PRJNA844206. The reference genome for *E. coli* GCF_000005845.2 is provided at https://www.ncbi.nlm.nih.gov/assembly/GCF_000005845.2, and for *V. cholerae* F_003063785.1 at https://www.ncbi.nlm.nih.gov/assembly/GCF_003063785.1. For *S. cerevisiae*, the reference genome of the W303 strain was used.

All other data supporting the findings of this study are available from the corresponding author on reasonable request. Source data are provided with this paper.

## Code availability

The custom-made code of the analysis is available online https://github.com/koszullab/T7_promoter_analysis.

Open-access versions of the programs and pipeline used (Hicstuff) are available online on the github account of the Koszul lab Hicstuff (www.github.com/koszullab/hicstuff) version 3.0.1, Bowtie284 (version 2.3.4.1 available online at http://bowtie-bio.sourceforge.net/bowtie2/), SAMtools85 (version 1.9 available online at http://www.htslib.org/download/http://www.htslib.org/download/), Bedtools86 (version 2.29.1 available online at https://bedtools.readthedocs.io/en/latest/content/installation.html) and Cooler81 (versions 0.8.7–0.8.11 available online at https://cooler.readthedocs.io/en/latest/).

---

**Supplementary Table 1:** Positions of the detected borders with the different methods and from Lioy *et al.* 2018 (Lioy et al. 2018).

| **Liroy *et al.* 2018** | **DI 5kb - window 100kb** | **DI 2kb - window 50kb** | **DI 1kb - window 20kb** | **HiBD 1kb - window: 10, 15, 20, 25, 30kb** |
| --- | --- | --- | --- | --- |
| 40 | 50 | 36 | 36;89;112 | 36;89;112 |
| 145 | 145 | 130 | 144;189 | 128;156;189;204 |
| 220 | 225 | 230 | 231;290;381 | 226;269;305;315;326;343;364;379;437 |
| 440 | 445 | 446 | 446;465 | 445 |
| 510 | 665 | 526;590;662 | 527;658;753 | 552;581;608;619;629;658;689;708;753 |
| 760 | 780;880 | 782;848;876 | 840;913;948 | 779;808;821;836;911;946 |
| 980 | 980 | 976;1016 | 986;1051 | 999;1013.1086;1106 |
| 1145 | 1140 | 1140 | 1140 | 1145;1167 |
| 1195 |  |  | 1245 | 1196;1268;1273;1300 |
| 1300 | 1315 | 1316;1402;1418 | 1317;1409;1471;1566 | 1318;1342;1366;1422;1437;1455;1497;1544;1549 |
| 1585 |  | 1598 | 1605 | 1601;1616;1621;1645 |
| 1670 | 1675 | 1702;1722 | 1674 | 1673;1702;1752;1757;1774 |
| 1795 | 1800 | 1800 | 1793 | 1789 |
| 1845 | 1970 | 1910;1970;2002 | 1843;1911;2970;2002;2024 | 1844;1865;1870;1906;1942;1963;2000;2024;2046;2084 |
| 2100 |  | 2108;2184 | 2139;2175;2228 | 2107;2130;2168 |
| 2255 | 2275 | 2270;2330 | 2304;2330;2369 | 2263 |
| 2380 | 2390 | 2390 | 2389;2425;2452 | 2389;2410;2425;2248;2495 |
| 2525 |  | 2538;2630 | 2540;2582;2611;2631;2682 | 2527;2537;2555;2614;2631;2691 |
| 2720 | 2730 | 2728 | 2730;2845;2886 | 2725;2756;2773;2811;2816;2833 |
| 2895 | 2905 | 2906;2952 |  | 2906;2942;2958 |
| 2990 | 3000 | 2988;3046;3062;3140 | 2982;3022;3066;3140 | 2973;3030;3156 |
| 3165 |  | 3194 | 3232 | 3222 |
| 3295 | 3300 | 3304;3360 | 3376;3392;3424 | 3304;3393;3425 |
| 3430 | 3440;3565 | 3442;3566 | 3446;3511 | 3440;3469;3511;3535;3566;3598 |
| 3645 |  | 3670 | 3701;3778 | 3635;3654;3696;3748 |
| 3800 | 3805 | 3808 | 3865;3910 | 3836;3876;3896;3910 |
| 3935 | 3945 | 3946 | 3948;3983 | 3943;3982;4004 |
| 4030 | 4045 | 4042 | 4041;4072;4146 | 4038;4062;4115;4136 |
| 4160 |  | 4170 | 4171;4190 | 4169;4190 |
| 4205 | 4210;4360 | 4212;4308;4362;4412 | 4214;4324;4386;4417 | 4210;4282;4385;4409 |
| 4465 | 4475 | 4474;4584 | 4475;4571 | 4473;4538;4458;4605 |

**Supplementary Table 2:**
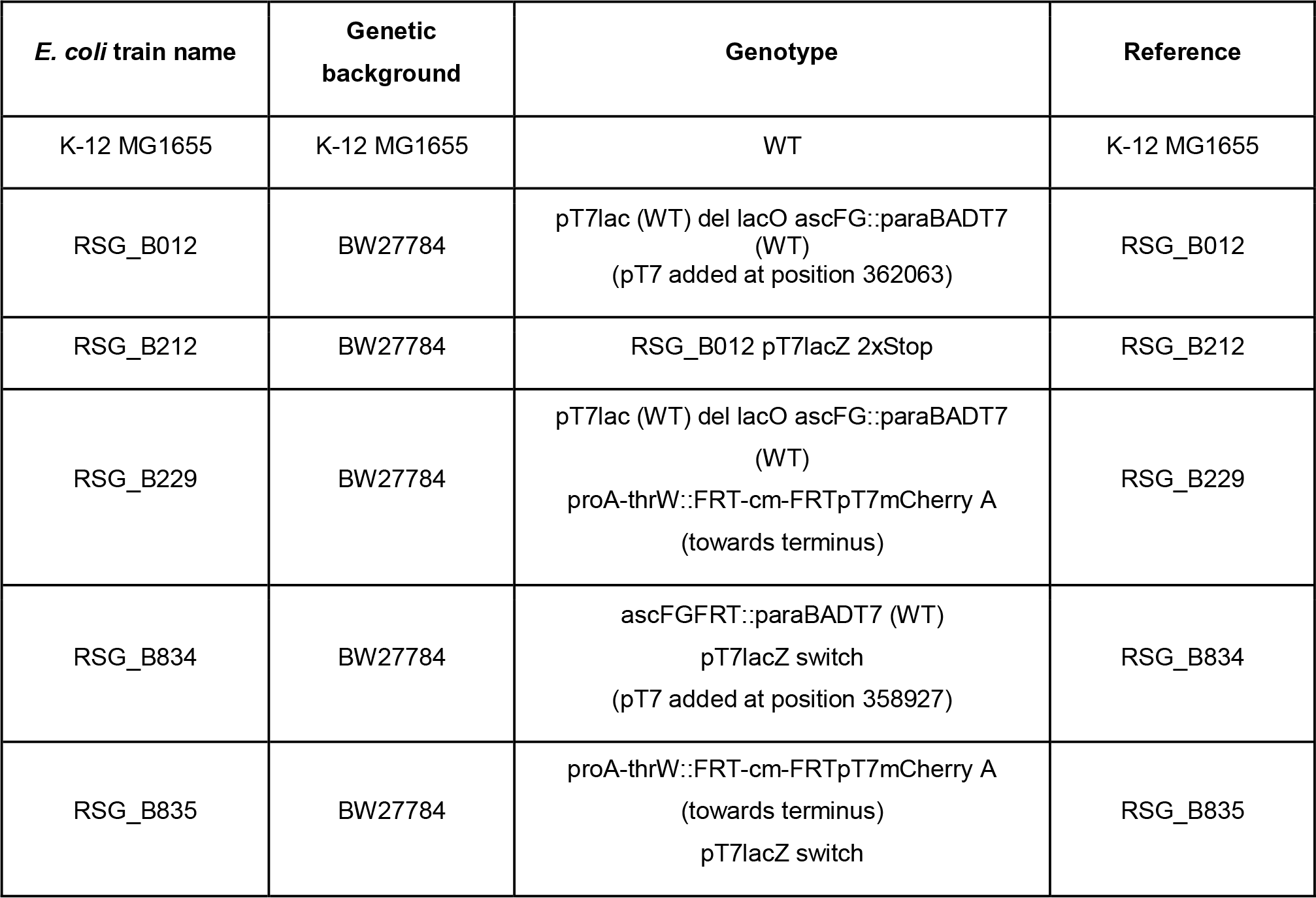
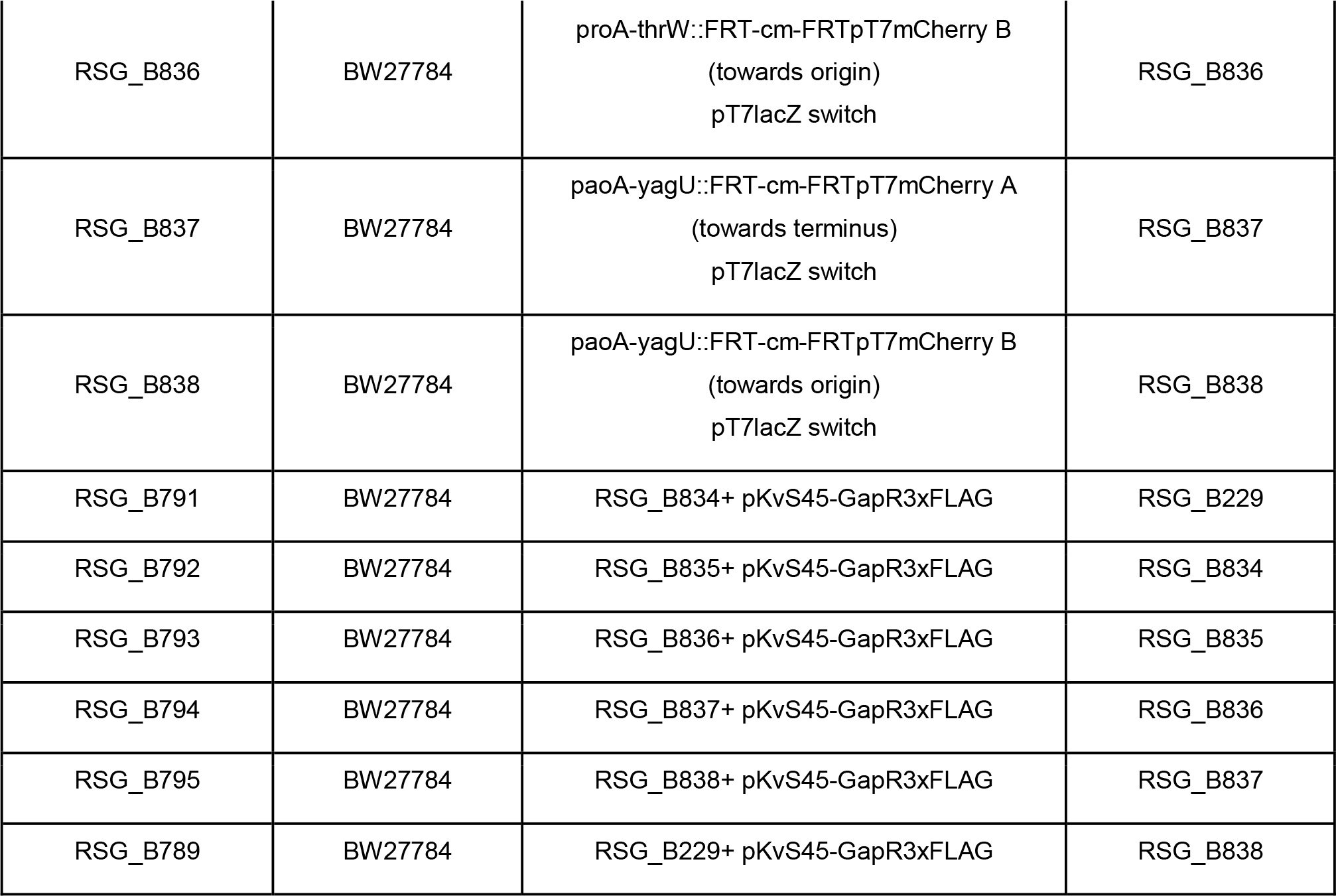
strains used in the study.

**Supplementary Table 3:**
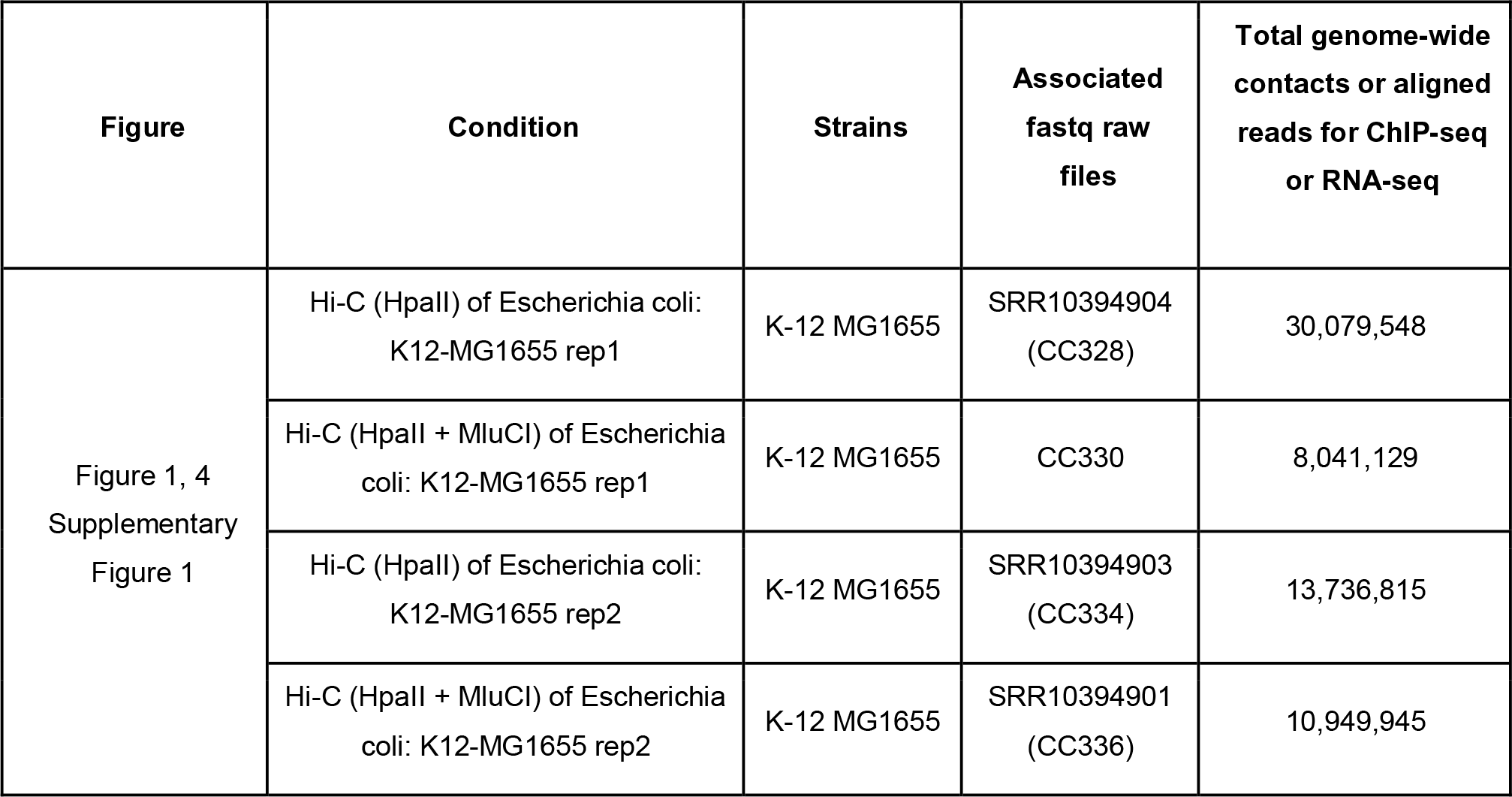
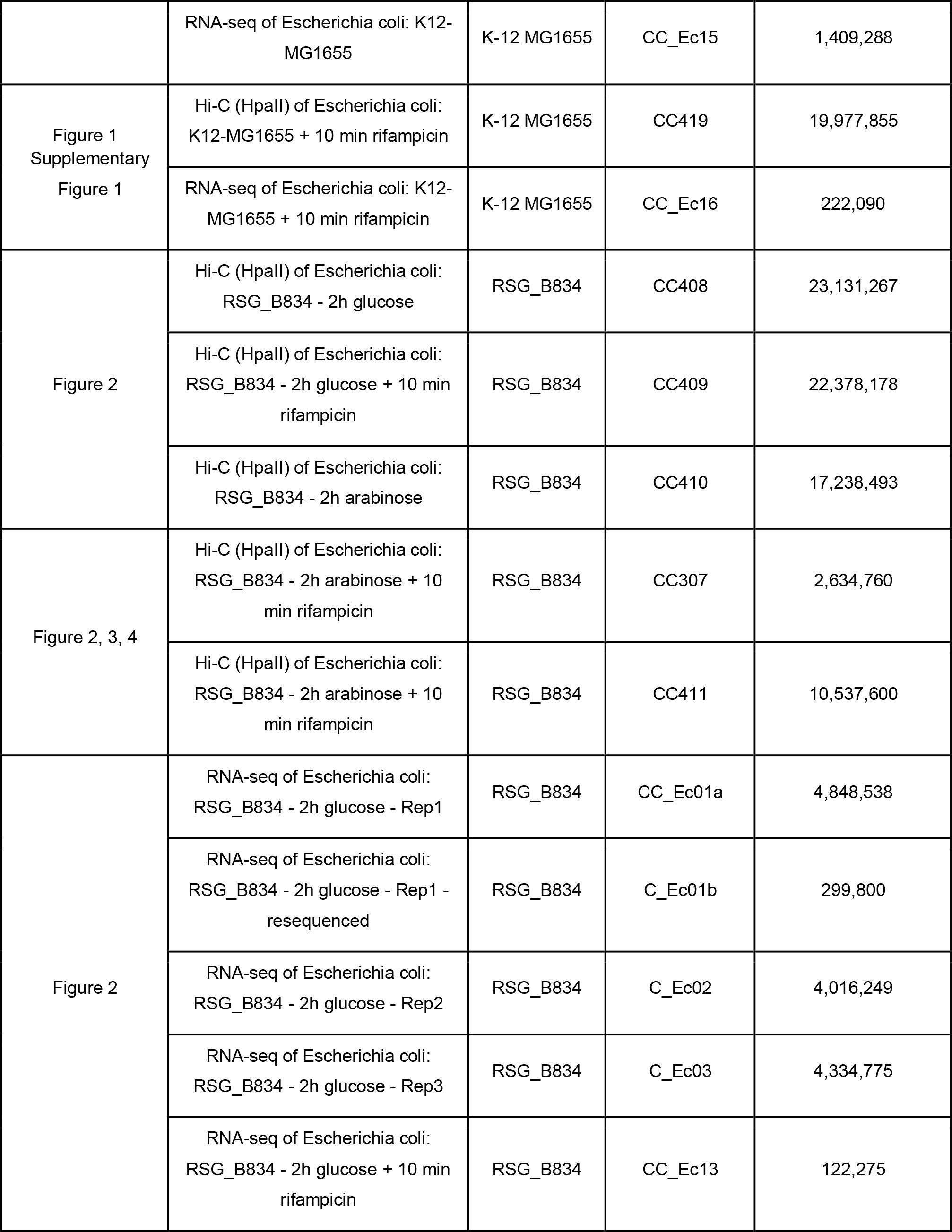
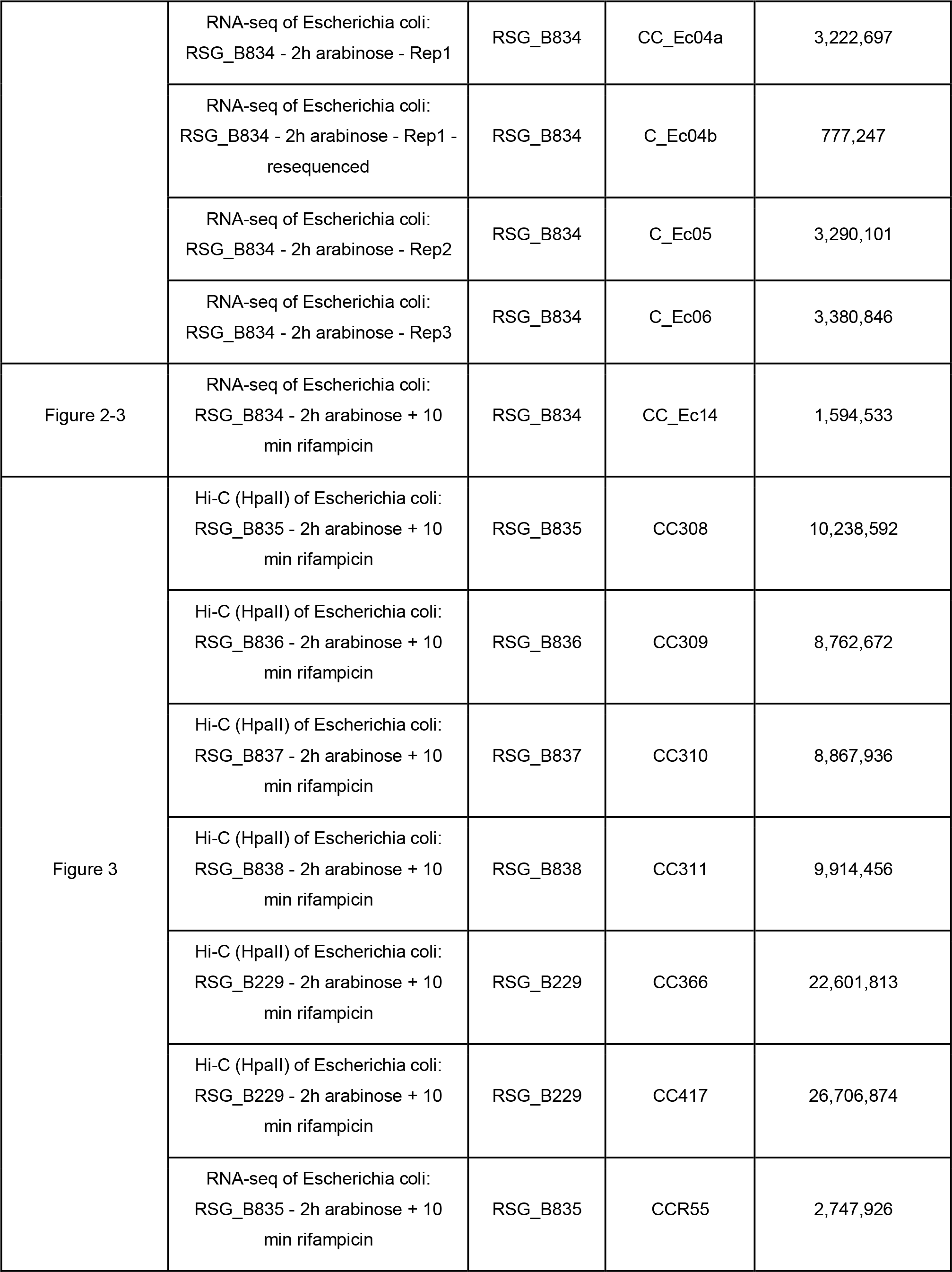
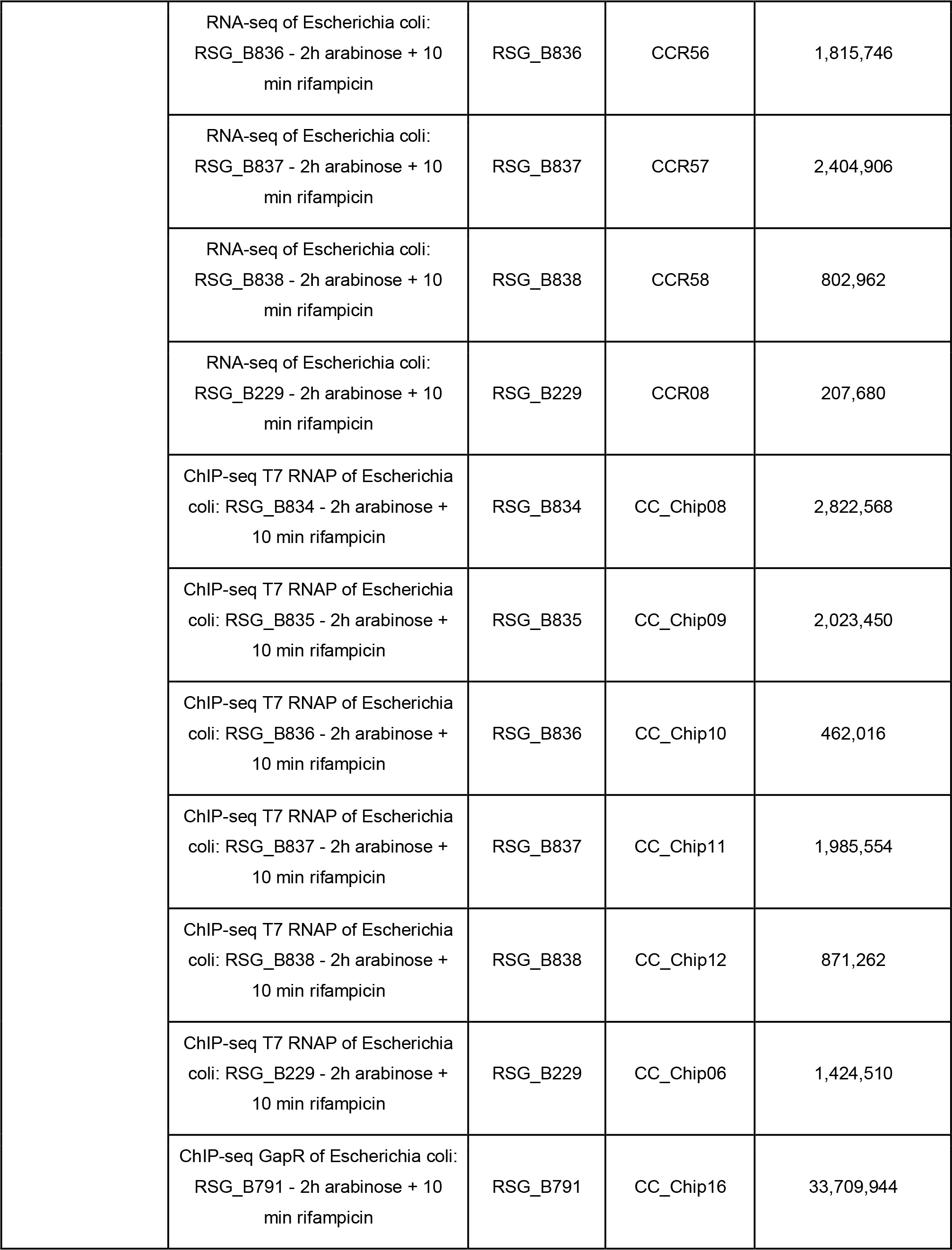
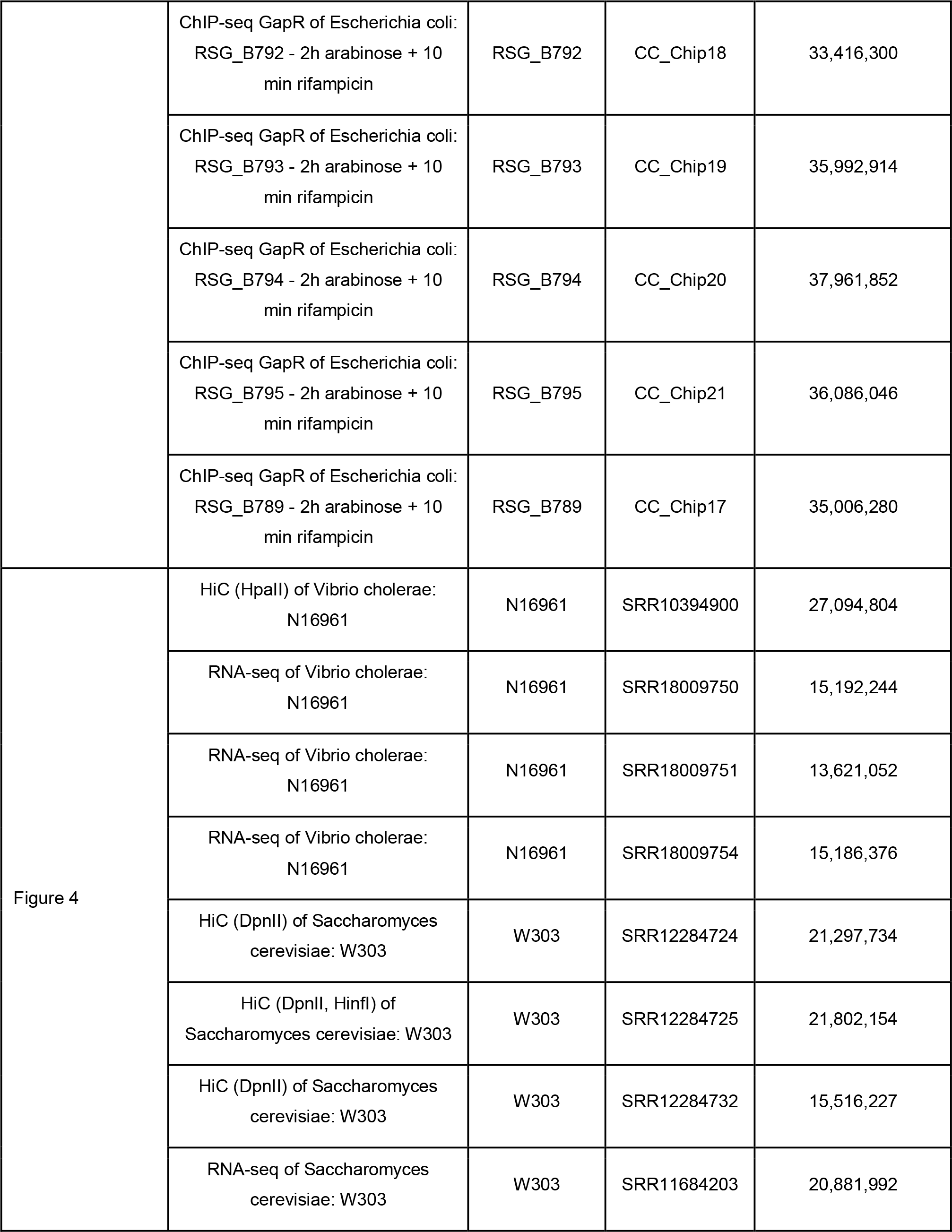
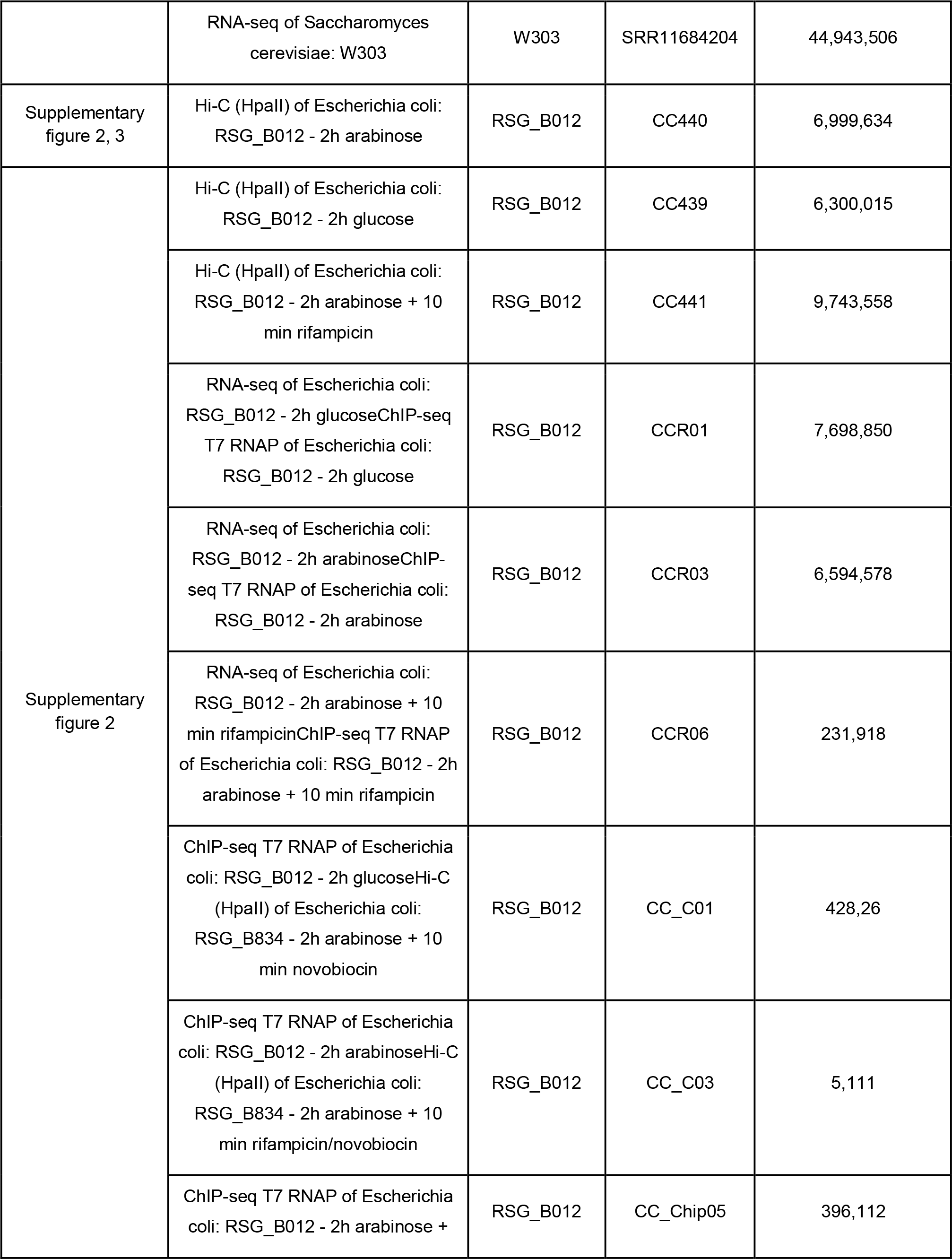
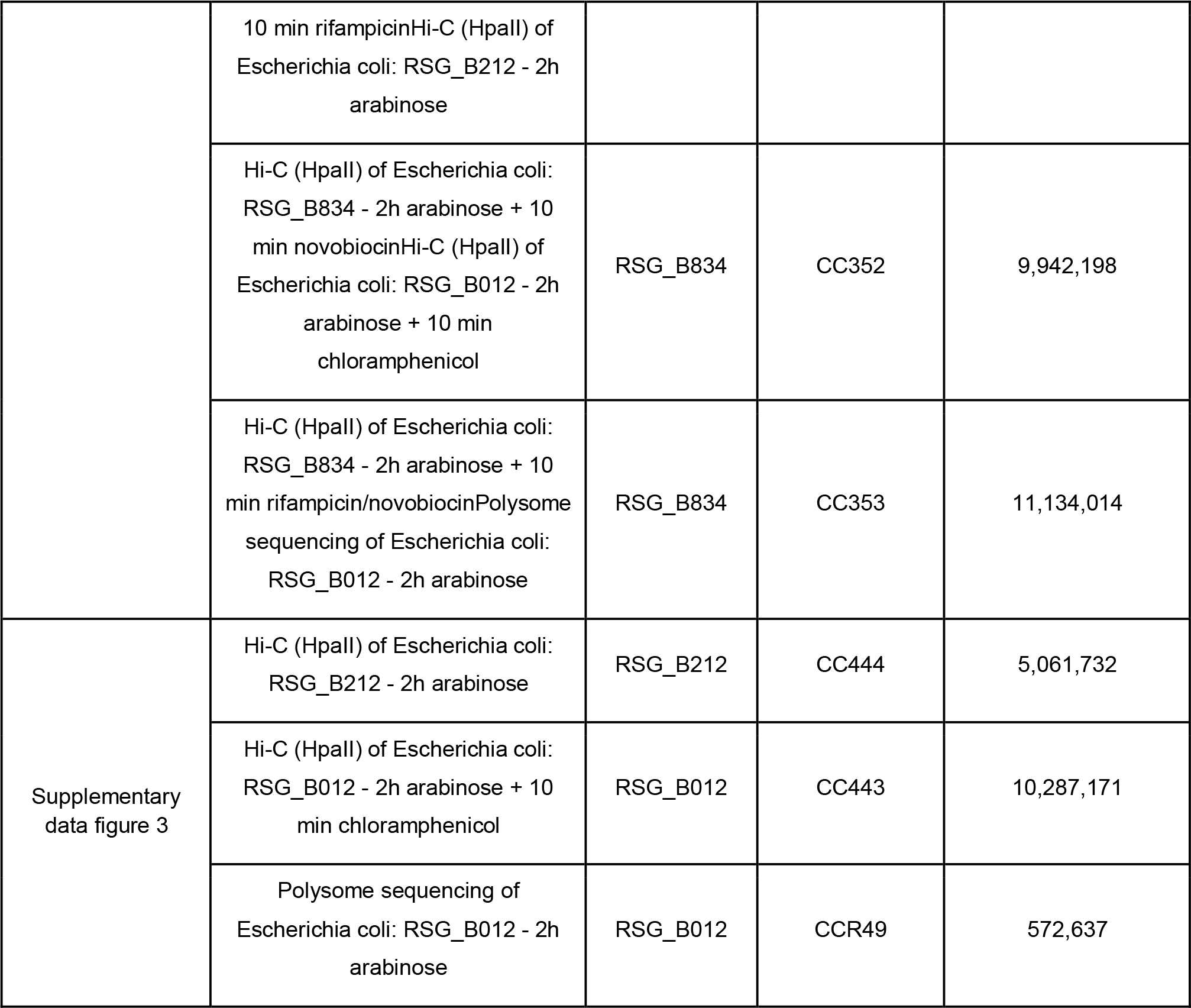
Libraries used in this study available at PRJNA844206.

| Figure | Condition | Strains | Associated fastq raw files | Total genome-wide contacts or aligned reads for ChIP-seq or RNA-seq |
| --- | --- | --- | --- | --- |
| Figure 1, 4<br>Supplementary Figure 1 | Hi-C (HpaII) of Escherichia coli: K12-MG1655 rep1 | K-12 MG1655 | SRR10394904 (CC328) | 30,079,548 |
|  | Hi-C (HpaII + MluCI) of Escherichia coli: K12-MG1655 rep1 | K-12 MG1655 | CC330 | 8,041,129 |
|  | Hi-C (HpaII) of Escherichia coli: K12-MG1655 rep2 | K-12 MG1655 | SRR10394903 (CC334) | 13,736,815 |
|  | Hi-C (HpaII + MluCI) of Escherichia coli: K12-MG1655 rep2 | K-12 MG1655 | SRR10394901 (CC336) | 10,949,945 |
|  | RNA-seq of Escherichia coli: K12-MG1655 | K-12 MG1655 | CC_Ec15 | 1,409,288 |
| Figure 1<br>Supplementary<br>Figure 1 | Hi-C (HpalI) of Escherichia coli: K12-MG1655 + 10 min rifampicin | K-12 MG1655 | CC419 | 19,977,855 |
|  | RNA-seq of Escherichia coli: K12-MG1655 + 10 min rifampicin | K-12 MG1655 | CC_Ec16 | 222,090 |
| Figure 2 | Hi-C (HpalI) of Escherichia coli: RSG_B834 - 2h glucose | RSG_B834 | CC408 | 23,131,267 |
|  | Hi-C (HpalI) of Escherichia coli: RSG_B834 - 2h glucose + 10 min rifampicin | RSG_B834 | CC409 | 22,378,178 |
|  | Hi-C (HpalI) of Escherichia coli: RSG_B834 - 2h arabinose | RSG_B834 | CC410 | 17,238,493 |
| Figure 2, 3, 4 | Hi-C (HpalI) of Escherichia coli: RSG_B834 - 2h arabinose + 10 min rifampicin | RSG_B834 | CC307 | 2,634,760 |
|  | Hi-C (HpalI) of Escherichia coli: RSG_B834 - 2h arabinose + 10 min rifampicin | RSG_B834 | CC411 | 10,537,600 |
| Figure 2 | RNA-seq of Escherichia coli: RSG_B834 - 2h glucose - Rep1 | RSG_B834 | CC_Ec01a | 4,848,538 |
|  | RNA-seq of Escherichia coli: RSG_B834 - 2h glucose - Rep1 - resequenced | RSG_B834 | C_Ec01b | 299,800 |
|  | RNA-seq of Escherichia coli: RSG_B834 - 2h glucose - Rep2 | RSG_B834 | C_Ec02 | 4,016,249 |
|  | RNA-seq of Escherichia coli: RSG_B834 - 2h glucose - Rep3 | RSG_B834 | C_Ec03 | 4,334,775 |
|  | RNA-seq of Escherichia coli: RSG_B834 - 2h glucose + 10 min rifampicin | RSG_B834 | CC_Ec13 | 122,275 |
|  | RNA-seq of Escherichia coli:<br>RSG_B834 - 2h arabinose - Rep1 | RSG_B834 | CC_Ec04a | 3,222,697 |
|  | RNA-seq of Escherichia coli:<br>RSG_B834 - 2h arabinose - Rep1 -<br>resequenced | RSG_B834 | C_Ec04b | 777,247 |
|  | RNA-seq of Escherichia coli:<br>RSG_B834 - 2h arabinose - Rep2 | RSG_B834 | C_Ec05 | 3,290,101 |
|  | RNA-seq of Escherichia coli:<br>RSG_B834 - 2h arabinose - Rep3 | RSG_B834 | C_Ec06 | 3,380,846 |
| Figure 2-3 | RNA-seq of Escherichia coli:<br>RSG_B834 - 2h arabinose + 10<br>min rifampicin | RSG_B834 | CC_Ec14 | 1,594,533 |
| Figure 3 | Hi-C (HpaII) of Escherichia coli:<br>RSG_B835 - 2h arabinose + 10<br>min rifampicin | RSG_B835 | CC308 | 10,238,592 |
|  | Hi-C (HpaII) of Escherichia coli:<br>RSG_B836 - 2h arabinose + 10<br>min rifampicin | RSG_B836 | CC309 | 8,762,672 |
|  | Hi-C (HpaII) of Escherichia coli:<br>RSG_B837 - 2h arabinose + 10<br>min rifampicin | RSG_B837 | CC310 | 8,867,936 |
|  | Hi-C (HpaII) of Escherichia coli:<br>RSG_B838 - 2h arabinose + 10<br>min rifampicin | RSG_B838 | CC311 | 9,914,456 |
|  | Hi-C (HpaII) of Escherichia coli:<br>RSG_B229 - 2h arabinose + 10<br>min rifampicin | RSG_B229 | CC366 | 22,601,813 |
|  | Hi-C (HpaII) of Escherichia coli:<br>RSG_B229 - 2h arabinose + 10<br>min rifampicin | RSG_B229 | CC417 | 26,706,874 |
|  | RNA-seq of Escherichia coli:<br>RSG_B835 - 2h arabinose + 10<br>min rifampicin | RSG_B835 | CCR55 | 2,747,926 |
|  | RNA-seq of Escherichia coli:<br>RSG_B836 - 2h arabinose + 10<br>min rifampicin | RSG_B836 | CCR56 | 1,815,746 |
|  | RNA-seq of Escherichia coli:<br>RSG_B837 - 2h arabinose + 10<br>min rifampicin | RSG_B837 | CCR57 | 2,404,906 |
|  | RNA-seq of Escherichia coli:<br>RSG_B838 - 2h arabinose + 10<br>min rifampicin | RSG_B838 | CCR58 | 802,962 |
|  | RNA-seq of Escherichia coli:<br>RSG_B229 - 2h arabinose + 10<br>min rifampicin | RSG_B229 | CCR08 | 207,680 |
|  | ChIP-seq T7 RNAP of Escherichia<br>coli: RSG_B834 - 2h arabinose +<br>10 min rifampicin | RSG_B834 | CC_Chip08 | 2,822,568 |
|  | ChIP-seq T7 RNAP of Escherichia<br>coli: RSG_B835 - 2h arabinose +<br>10 min rifampicin | RSG_B835 | CC_Chip09 | 2,023,450 |
|  | ChIP-seq T7 RNAP of Escherichia<br>coli: RSG_B836 - 2h arabinose +<br>10 min rifampicin | RSG_B836 | CC_Chip10 | 462,016 |
|  | ChIP-seq T7 RNAP of Escherichia<br>coli: RSG_B837 - 2h arabinose +<br>10 min rifampicin | RSG_B837 | CC_Chip11 | 1,985,554 |
|  | ChIP-seq T7 RNAP of Escherichia<br>coli: RSG_B838 - 2h arabinose +<br>10 min rifampicin | RSG_B838 | CC_Chip12 | 871,262 |
|  | ChIP-seq T7 RNAP of Escherichia<br>coli: RSG_B229 - 2h arabinose +<br>10 min rifampicin | RSG_B229 | CC_Chip06 | 1,424,510 |
|  | ChIP-seq GapR of Escherichia coli:<br>RSG_B791 - 2h arabinose + 10<br>min rifampicin | RSG_B791 | CC_Chip16 | 33,709,944 |
|  | ChIP-seq GapR of Escherichia coli:<br>RSG_B792 - 2h arabinose + 10<br>min rifampicin | RSG_B792 | CC_Chip18 | 33,416,300 |
|  | ChIP-seq GapR of Escherichia coli:<br>RSG_B793 - 2h arabinose + 10<br>min rifampicin | RSG_B793 | CC_Chip19 | 35,992,914 |
|  | ChIP-seq GapR of Escherichia coli:<br>RSG_B794 - 2h arabinose + 10<br>min rifampicin | RSG_B794 | CC_Chip20 | 37,961,852 |
|  | ChIP-seq GapR of Escherichia coli:<br>RSG_B795 - 2h arabinose + 10<br>min rifampicin | RSG_B795 | CC_Chip21 | 36,086,046 |
|  | ChIP-seq GapR of Escherichia coli:<br>RSG_B789 - 2h arabinose + 10<br>min rifampicin | RSG_B789 | CC_Chip17 | 35,006,280 |
| Figure 4 | HiC (HpaII) of Vibrio cholerae:<br>N16961 | N16961 | SRR10394900 | 27,094,804 |
|  | RNA-seq of Vibrio cholerae:<br>N16961 | N16961 | SRR18009750 | 15,192,244 |
|  | RNA-seq of Vibrio cholerae:<br>N16961 | N16961 | SRR18009751 | 13,621,052 |
|  | RNA-seq of Vibrio cholerae:<br>N16961 | N16961 | SRR18009754 | 15,186,376 |
|  | HiC (DpnII) of Saccharomyces<br>cerevisiae: W303 | W303 | SRR12284724 | 21,297,734 |
|  | HiC (DpnII, HinfI) of<br>Saccharomyces cerevisiae: W303 | W303 | SRR12284725 | 21,802,154 |
|  | HiC (DpnII) of Saccharomyces<br>cerevisiae: W303 | W303 | SRR12284732 | 15,516,227 |
|  | RNA-seq of Saccharomyces<br>cerevisiae: W303 | W303 | SRR11684203 | 20,881,992 |
|  | RNA-seq of *Saccharomyces cerevisiae*: W303 | W303 | SRR11684204 | 44,943,506 |
| Supplementary figure 2, 3 | Hi-C (HpalI) of *Escherichia coli*: RSG_B012 - 2h arabinose | RSG_B012 | CC440 | 6,999,634 |
| Supplementary figure 2 | Hi-C (HpalI) of *Escherichia coli*: RSG_B012 - 2h glucose | RSG_B012 | CC439 | 6,300,015 |
|  | Hi-C (HpalI) of *Escherichia coli*: RSG_B012 - 2h arabinose + 10 min rifampicin | RSG_B012 | CC441 | 9,743,558 |
|  | RNA-seq of *Escherichia coli*: RSG_B012 - 2h glucoseChIP-seq T7 RNAP of *Escherichia coli*: RSG_B012 - 2h glucose | RSG_B012 | CCR01 | 7,698,850 |
|  | RNA-seq of *Escherichia coli*: RSG_B012 - 2h arabinoseChIP-seq T7 RNAP of *Escherichia coli*: RSG_B012 - 2h arabinose | RSG_B012 | CCR03 | 6,594,578 |
|  | RNA-seq of *Escherichia coli*: RSG_B012 - 2h arabinose + 10 min rifampicinChIP-seq T7 RNAP of *Escherichia coli*: RSG_B012 - 2h arabinose + 10 min rifampicin | RSG_B012 | CCR06 | 231,918 |
|  | ChIP-seq T7 RNAP of *Escherichia coli*: RSG_B012 - 2h glucoseHi-C (HpalI) of *Escherichia coli*: RSG_B834 - 2h arabinose + 10 min novobiocin | RSG_B012 | CC_C01 | 428,26 |
|  | ChIP-seq T7 RNAP of *Escherichia coli*: RSG_B012 - 2h arabinoseHi-C (HpalI) of *Escherichia coli*: RSG_B834 - 2h arabinose + 10 min rifampicin/novobiocin | RSG_B012 | CC_C03 | 5,111 |
|  | ChIP-seq T7 RNAP of *Escherichia coli*: RSG_B012 - 2h arabinose + | RSG_B012 | CC_Chip05 | 396,112 |
|  | 10 min rifampicinHi-C (HpalI) of Escherichia coli: RSG_B212 - 2h arabinose |  |  |  |
|  | Hi-C (HpalI) of Escherichia coli: RSG_B834 - 2h arabinose + 10 min novobiocinHi-C (HpalI) of Escherichia coli: RSG_B012 - 2h arabinose + 10 min chloramphenicol | RSG_B834 | CC352 | 9,942,198 |
|  | Hi-C (HpalI) of Escherichia coli: RSG_B834 - 2h arabinose + 10 min rifampicin/novobiocinPolysome sequencing of Escherichia coli: RSG_B012 - 2h arabinose | RSG_B834 | CC353 | 11,134,014 |
| Supplementary data figure 3 | Hi-C (HpalI) of Escherichia coli: RSG_B212 - 2h arabinose | RSG_B212 | CC444 | 5,061,732 |
|  | Hi-C (HpalI) of Escherichia coli: RSG_B012 - 2h arabinose + 10 min chloramphenicol | RSG_B012 | CC443 | 10,287,171 |
|  | Polysome sequencing of Escherichia coli: RSG_B012 - 2h arabinose | RSG_B012 | CCR49 | 572,637 |

## Supplementary Figures

**Supplementary Data Figure 1:**
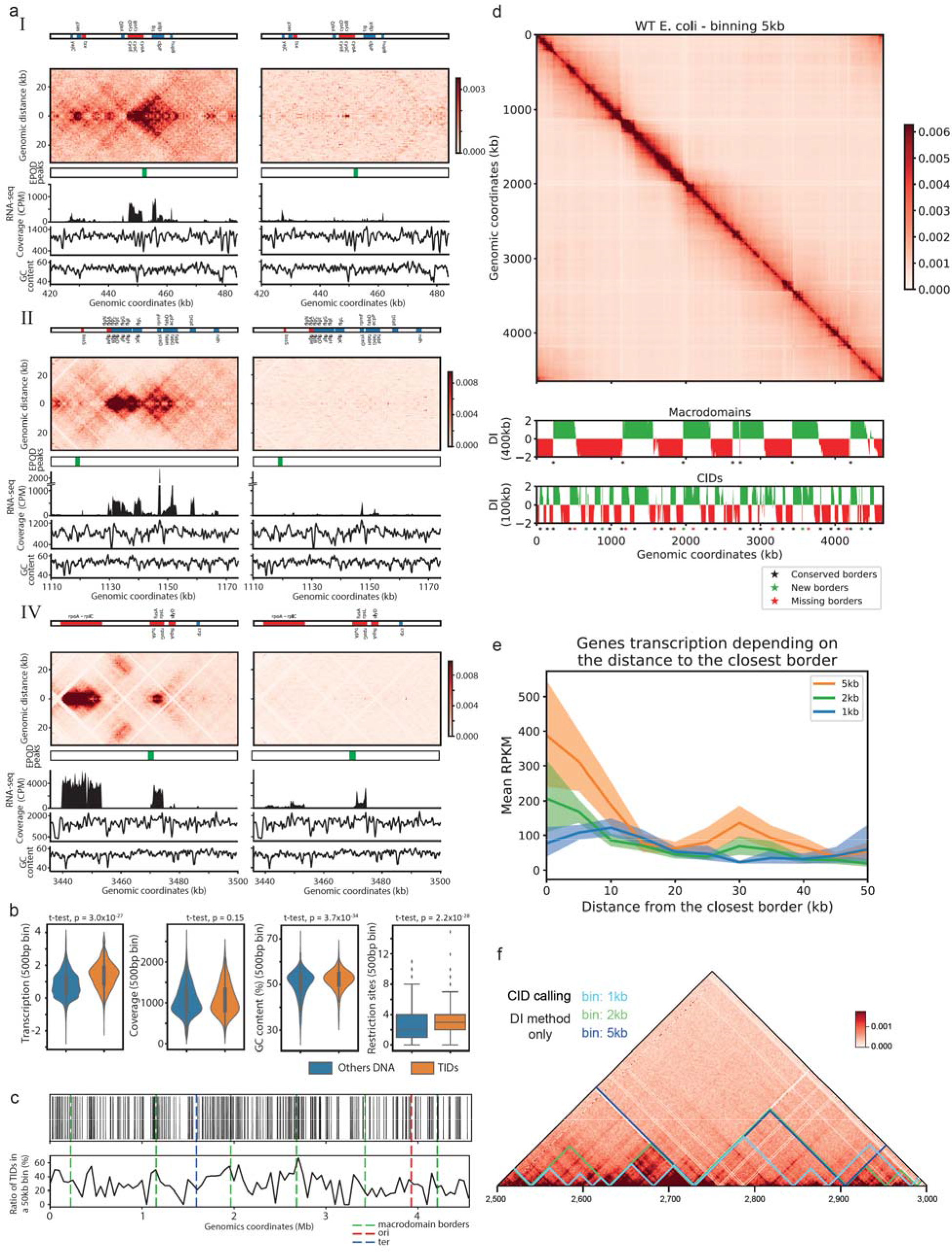
Transcription impact on WT bacterial chromosome folding. **a,** Magnifications of regions I, II and IV. The names of the genes within the 10% most transcribed are indicated (blue and red correspond to forwards and reverse genes, respectively). Left panels: normalized contact map (bin: 0.5 kb) over the corresponding EPODs peaks11 and RNA-seq profile (in Count Per Million or CPM), Hi-C coverage, GC content (%), in absence of rifampicin. Right panel: same region and analysis but in presence of rifampicin. **b,** From left to right: distributions of transcription (CPM, in log 10), coverage, GC content and numbers of restrictions sites in pairs of bins with either low (blue) and high (i.e. in TIDs; orange) contact frequency at short range (Methods). The p-values are from independent t-tests. **c,** Distributions of the local thick bundle across the whole genome (x axis). Top: each strip represents a 500 bp bin called within a thick bundle (i.e. TID; Methods). Bottom: same data as above but binned into 50 kb bins. The positions of the macrodomains as defined in Lioy et al. (Lioy et al. 2018) are indicated by green dotted lines. Ori and ter are indicted by red and blue lines, respectively. **d,** *E coli* contact map binned at 5kb at the top. Below the corresponding detected macrodomains and CIDs based on directional index method7. Stars show the significant borders detected, in both Lioy *et al.* data and our data (black), only detected in Lioy et al. data (red) and only detected with our data (green). **e,** Gene transcription in RPKM depending on the distance from the closest borders detected at different resolutions. Error intervals are generated from bootstraps. **f,** Domains detected based on DI analysis only at different resolutions; 1kb (cyan), 2kb (green) and 5kb (blue).

**Supplementary Data Figure 2:**
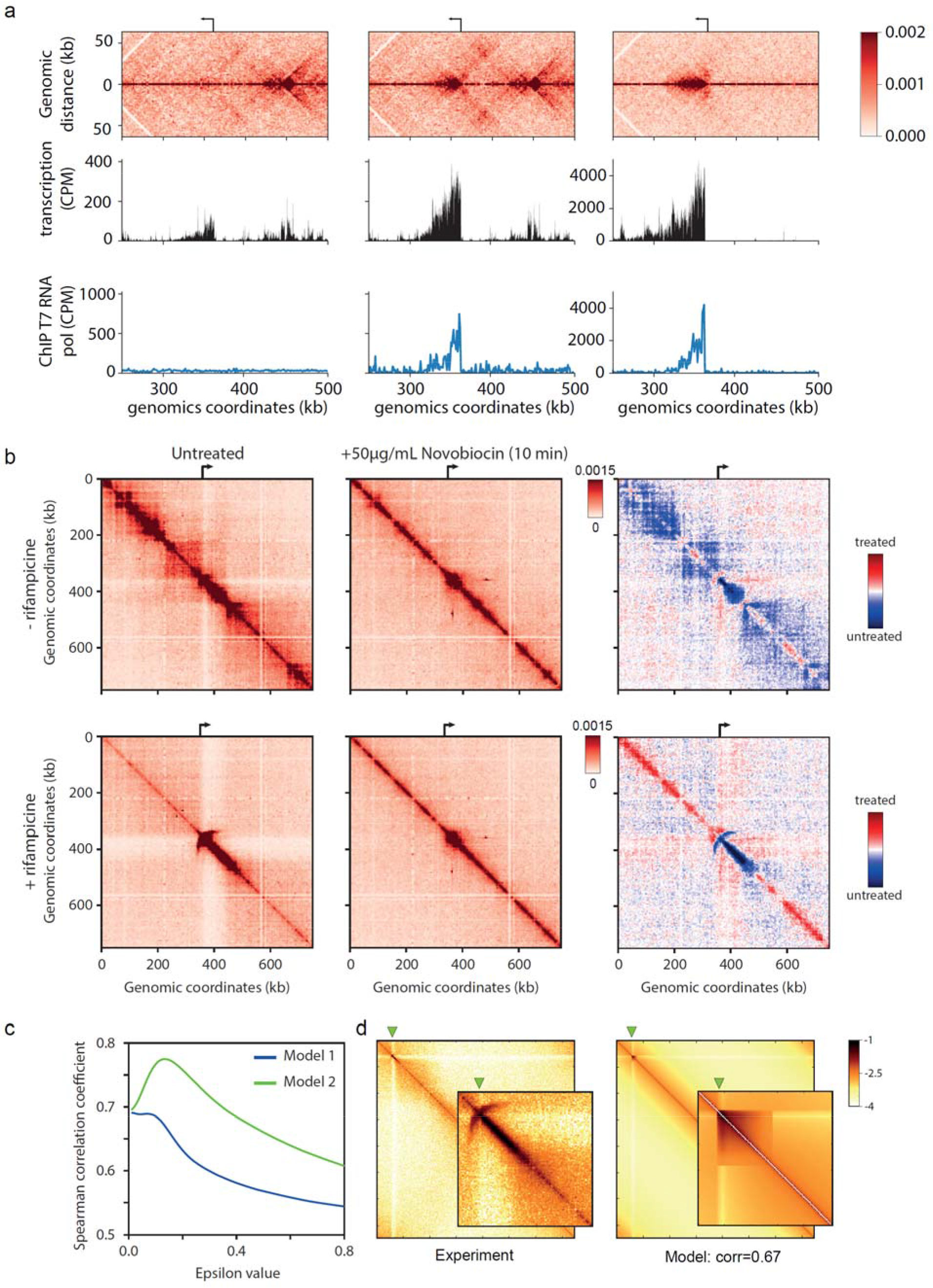
Activation of a single transcription unit within the E. coli chromosome. **a**, Magnifications of the Hi-C contact maps (bin: 1kb) of *E. coli* chromosome carrying a single T7 promoter facing toward the ori, with below the corresponding RNAseq and the signal from ChIP of the T7 RNA polymerase. From left to right: the T7 promoter off, the T7 promoter on and the T7 promoter on with rifampicin. **b**, Hi-C contact map magnifications (bin: 1kb) of the bacteria carrying T7 promoter facing to the terminus. From left to right: without novobiocin treatment, with novobiocin treatment, log2 ratio of treated over untreated contact maps. In bottom contact rifampicin have been added. **c,** Correlation between the maps recovered from each of the two models and the experimental map, depending on the epsilon values (Methods). **d**, Best correlation map of Model I (right), aside the experimental map (left).

**Supplementary Data Figure 3:**
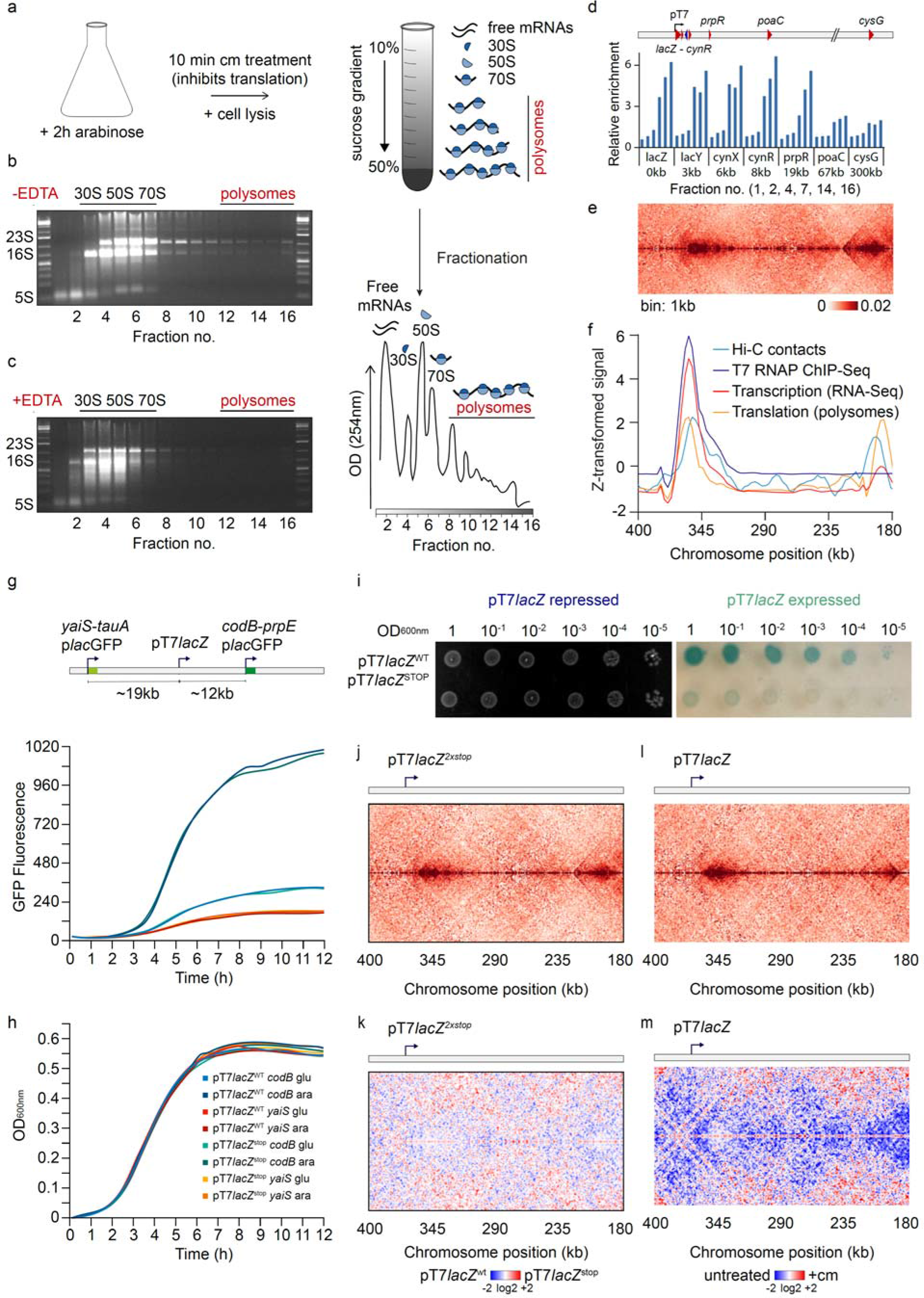
Translation impact on bacterial chromosome folding. **a**, Schematic view of the polysome extraction experiment. **b, c**, Gel migration of the different fractions for polysome extraction without EDTA (**b**), and with EDTA (**c**). **d**, Magnification of the Hi-C contact map of the *E. coli* carrying T7 promoter facing the origin. **e**, Corresponding z-transformed signals of the short range Hi-C signal, T7 RNA polymerase ChIP-seq, transcription and translation. **f, g**, Gene expression upstream (yaiS) and downstream (codB) of the T7 promoter lacZ system with or without STOP codons based on GFP fluorescence (**f**) and growth of the corresponding strains (**g**). **h**, Bacterial colony dilution with pT7lacZ repressed on the left and expressed on the right. **i**, Contact map of the bacteria carrying a T7 promoter lacZ system with two stop codons into the lacZ gene. **j**, Log2 ratio between the contact map with the lacZ2xSTOP system over the contact map with the WT lacZ. **k**, Contact map of the bacteria carrying a T7 promoter lacZ system treated with chloramphenicol. **l**, Log2 ratio between the contact map treated over the untreated.

**Supplementary Data Figure 4:**
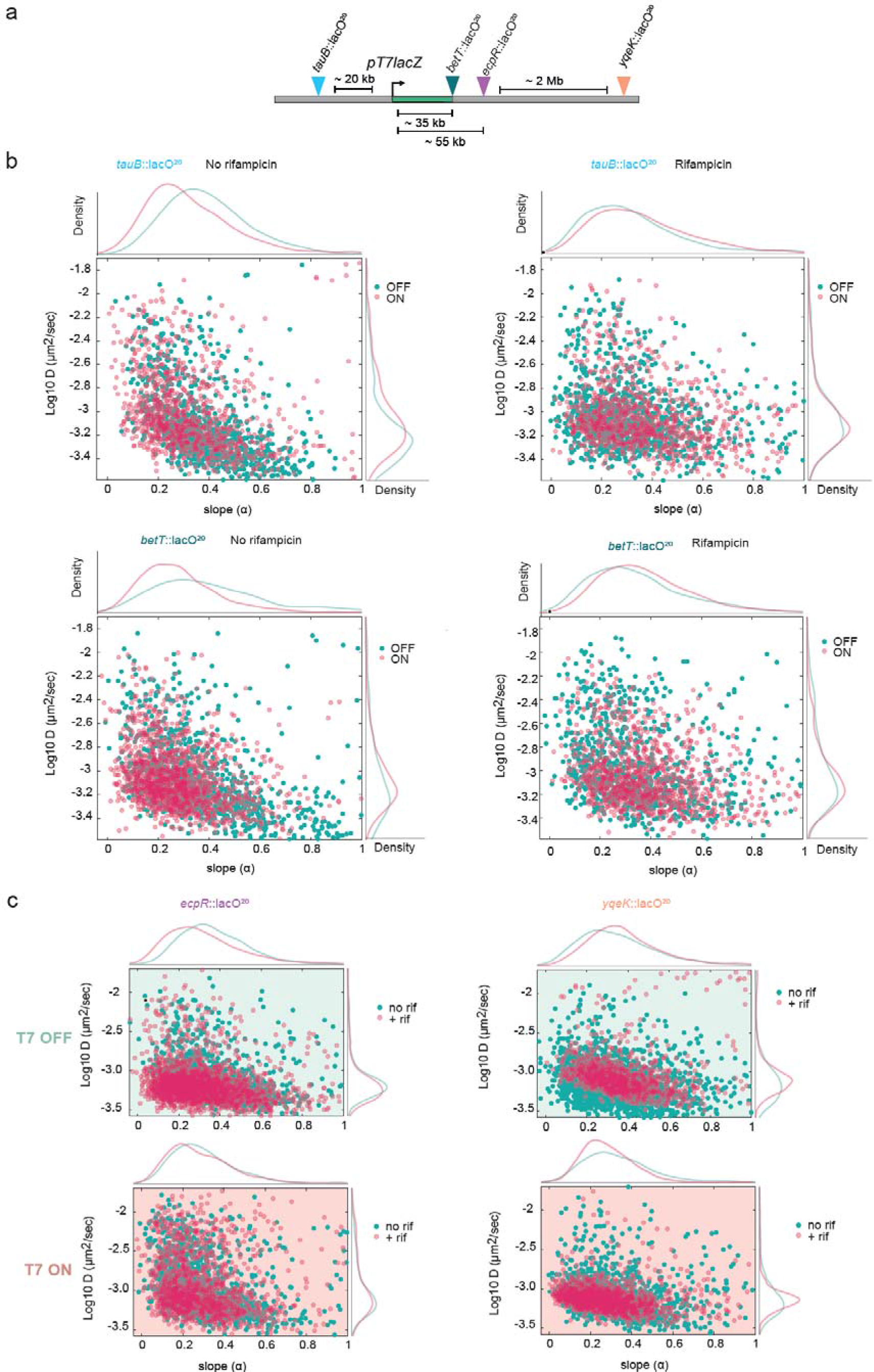
Dynamics of the T7 transcription unit. **a,** Positions of the four *lacO* arrays inserted in the vicinity of the T7 promoter. **b,** Scatter plots of D_c_ in function of the MSD slope for ~1000 trajectories upon induction of the T7 TU in the absence or presence of rifampicin. Top panels: the *tauB* locus at 20kb upstream of T7 TU TSS. Bottom panels: the *betT* locus at 35 kb downstream from T7 TU TSS. The mean (μ) of each distribution is indicated, their statistical differences measured by an Anova Kruskal-Wallis test with Bonferroni correction, * <0.033, ** < 0.0021, **** <0.0002, ****<0.0001. **c,** Comparison of *lacO* dynamics for the *ecpR* locus (left) and the *yqeK* locus (right) according to the presence of rifampicin; top in the absence of T7 transcription; bottom in the presence of T7 transcription.

## Notes

### Competing Interest Statement

The authors have declared no competing interest.

## References

Banigan EJ, Tang W, van den Berg AA, Stocsits RR, Wutz G, Brandão HB, Busslinger GA, Peters J-M, Mirny LA. 2022. Transcription shapes 3D chromatin organization by interacting with loop-extruding cohesin complexes. Genomics http://biorxiv.org/lookup/doi/10.1101/2022.01.07.475367 (Accessed June 3, 2022).

Baudry L, Millot GA, Thierry A, Koszul R, Scolari VF. 2020. Serpentine: a flexible 2D binning method for differential Hi-C analysis ed. A. Valencia. Bioinformatics 36: 3645–3651.

Brandão HB, Paul P, van den Berg AA, Rudner DZ, Wang X, Mirny LA. 2019. RNA polymerases as moving barriers to condensin loop extrusion. Proc Natl Acad Sci U S A 116: 20489–20499.

Chen F, Li G, Zhang MQ, Chen Y. 2018. HiCDB: a sensitive and robust method for detecting contact domain boundaries. Nucleic Acids Res 46: 11239–11250.

Cockram C, Thierry A, Gorlas A, Lestini R, Koszul R. 2021a. Euryarchaeal genomes are folded into SMC-dependent loops and domains, but lack transcription-mediated compartmentalization. Mol Cell 81: 459–472.e10.

Cockram C, Thierry A, Koszul R. 2021b. Generation of gene-level resolution chromosome contact maps in bacteria and archaea. STAR Protoc 2: 100512.

Cockram CA, Filatenkova M, Danos V, El Karoui M, Leach DRF. 2015. Quantitative genomic analysis of RecA protein binding during DNA double-strand break repair reveals RecBCD action in vivo. Proc Natl Acad Sci 112. https://pnas.org/doi/full/10.1073/pnas.1424269112 (Accessed June 3, 2022).

Cournac A, Marie-Nelly H, Marbouty M, Koszul R, Mozziconacci J. 2012. Normalization of a chromosomal contact map. BMC Genomics 13: 436.

Dame RT, Rashid F-ZM, Grainger DC. 2020. Chromosome organization in bacteria: mechanistic insights into genome structure and function. Nat Rev Genet 21: 227–242.

Deng S, Stein RA, Higgins NP. 2004. Transcription-induced barriers to supercoil diffusion in the *Salmonella typhimurium* chromosome. Proc Natl Acad Sci 101: 3398–3403.

Dixon JR, Selvaraj S, Yue F, Kim A, Li Y, Shen Y, Hu M, Liu JS, Ren B. 2012. Topological domains in mammalian genomes identified by analysis of chromatin interactions. Nature 485: 376–380.

Dorman CJ. 2019. DNA supercoiling and transcription in bacteria: a two-way street. BMC Mol Cell Biol 20: 26.

Freddolino PL, Amemiya HM, Goss TJ, Tavazoie S. 2021. Dynamic landscape of protein occupancy across the Escherichia coli chromosome ed. M.K. Waldor. PLOS Biol 19: e3001306.

Gaal T, Bratton BP, Sanchez-Vazquez P, Sliwicki A, Sliwicki K, Vegel A, Pannu R, Gourse RL. 2016. Colocalization of distant chromosomal loci in space in *E. coli*□: a bacterial nucleolus. Genes Dev 30: 2272–2285.

Ganji M, Shaltiel IA, Bisht S, Kim E, Kalichava A, Haering CH, Dekker C. 2018. Real-time imaging of DNA loop extrusion by condensin. Science 360: 102–105.

Germier T, Kocanova S, Walther N, Bancaud A, Shaban HA, Sellou H, Politi AZ, Ellenberg J, Gallardo F, Bystricky K. 2017. Real-Time Imaging of a Single Gene Reveals Transcription-Initiated Local Confinement. Biophys J 113: 1383–1394.

Gu B, Swigut T, Spencley A, Bauer MR, Chung M, Meyer T, Wysocka J. 2018. Transcription-coupled changes in nuclear mobility of mammalian cis-regulatory elements. Science 359: 1050–1055.

Guo MS, Kawamura R, Littlehale ML, Marko JF, Laub MT. 2021. High-resolution, genome-wide mapping of positive supercoiling in chromosomes. eLife 10: e67236.

Hsieh T-HS, Fudenberg G, Goloborodko A, Rando OJ. 2016. Micro-C XL: assaying chromosome conformation from the nucleosome to the entire genome. Nat Methods 13: 1009–1011.

Imakaev M, Fudenberg G, McCord RP, Naumova N, Goloborodko A, Lajoie BR, Dekker J, Mirny LA. 2012. Iterative correction of Hi-C data reveals hallmarks of chromosome organization. Nat Methods 9: 999–1003.

Kleckner N, Fisher JK, Stouf M, White MA, Bates D, Witz G. 2014. The bacterial nucleoid: nature, dynamics and sister segregation. Curr Opin Microbiol 22: 127–137.

Ladouceur A-M, Parmar BS, Biedzinski S, Wall J, Tope SG, Cohn D, Kim A, Soubry N, Reyes-Lamothe R, Weber SC. 2020. Clusters of bacterial RNA polymerase are biomolecular condensates that assemble through liquid-liquid phase separation. Proc Natl Acad Sci U S A 117: 18540–18549.

Le TB, Laub MT. 2016. Transcription rate and transcript length drive formation of chromosomal interaction domain boundaries. EMBO J 35: 1582–1595.

Le TBK, Imakaev MV, Mirny LA, Laub MT. 2013. High-resolution mapping of the spatial organization of a bacterial chromosome. Science 342: 731–734.

Lesne A, Riposo J, Roger P, Cournac A, Mozziconacci J. 2014. 3D genome reconstruction from chromosomal contacts. Nat Methods 11: 1141–1143.

Levine C, Hiasa H, Marians KJ. 1998. DNA gyrase and topoisomerase IV: biochemical activities, physiological roles during chromosome replication, and drug sensitivities. Biochim Biophys Acta 1400: 29–43.

Lioy VS, Cournac A, Marbouty M, Duigou S, Mozziconacci J, Espéli O, Boccard F, Koszul R. 2018. Multiscale Structuring of the E. coli Chromosome by Nucleoid-Associated and Condensin Proteins. Cell 172: 771–783.e18.

Lioy VS, Junier I, Boccard F. 2021. Multiscale Dynamic Structuring of Bacterial Chromosomes. Annu Rev Microbiol 75: 541–561.

Liu LF, Wang JC. 1987. Supercoiling of the DNA template during transcription. Proc Natl Acad Sci 84: 7024–7027.

Mäkelä J, Sherratt DJ. 2020. Organization of the Escherichia coli Chromosome by a MukBEF Axial Core. Mol Cell 78: 250–260.e5.

Marbouty M, Le Gall A, Cattoni DI, Cournac A, Koh A, Fiche J-B, Mozziconacci J, Murray H, Koszul R, Nollmann M. 2015. Condensin- and Replication-Mediated Bacterial Chromosome Folding and Origin Condensation Revealed by Hi-C and Super-resolution Imaging. Mol Cell 59: 588–602.

Matthey-Doret C, Baudry L, Bignaud A, Cournac A, Montagne R, Guiglielmoni N, Foutel-Rodier T, Scolari VF. 2020. koszullab/hicstuff: Use miniconda layer for docker and improved P(s) normalisation. Zenodo, October 5.

Postow L, Hardy CD, Arsuaga J, Cozzarelli NR. 2004. Topological domain structure of the Escherichia coli chromosome. Genes Dev 18: 1766–1779.

Racko D, Benedetti F, Dorier J, Stasiak A. 2018. Transcription-induced supercoiling as the driving force of chromatin loop extrusion during formation of TADs in interphase chromosomes. Nucleic Acids Res 46: 1648–1660.

Scolari VF, Mercy G, Koszul R, Lesne A, Mozziconacci J. 2018. Kinetic Signature of Cooperativity in the Irreversible Collapse of a Polymer. Phys Rev Lett 121: 057801.

Stracy M, Lesterlin C, Garza de Leon F, Uphoff S, Zawadzki P, Kapanidis AN. 2015. Live-cell superresolution microscopy reveals the organization of RNA polymerase in the bacterial nucleoid. Proc Natl Acad Sci 112. https://pnas.org/doi/full/10.1073/pnas.1507592112 (Accessed June 3, 2022).

Tabor S, Richardson CC. 1985. A bacteriophage T7 RNA polymerase/promoter system for controlled exclusive expression of specific genes. Proc Natl Acad Sci 82: 1074–1078.

ten Heggeler-Bordier B, Wahli W, Adrian M, Stasiak A, Dubochet J. 1992. The apical localization of transcribing RNA polymerases on supercoiled DNA prevents their rotation around the template. EMBO J 11: 667–672.

Umbarger MA, Toro E, Wright MA, Porreca GJ, Baù D, Hong S-H, Fero MJ, Zhu LJ, Marti-Renom MA, McAdams HH, et al. 2011. The Three-Dimensional Architecture of a Bacterial Genome and Its Alteration by Genetic Perturbation. Mol Cell 44: 252–264.

Valens M, Penaud S, Rossignol M, Cornet F, Boccard F. 2004. Macrodomain organization of the Escherichia coli chromosome. EMBO J 23: 4330–4341.

Vickridge E, Planchenault C, Cockram C, Junceda IG, Espéli O. 2017. Management of E. coli sister chromatid cohesion in response to genotoxic stress. Nat Commun 8: 14618.

Visser BJ, Sharma S, Chen PJ, McMullin AB, Bates ML, Bates D. 2022. Psoralen mapping reveals a bacterial genome supercoiling landscape dominated by transcription. Nucleic Acids Res 50: 4436–4449.

Wang X, Brandão HB, Le TBK, Laub MT, Rudner DZ. 2017. Bacillus subtilis SMC complexes juxtapose chromosome arms as they travel from origin to terminus. Science 355: 524–527.

Worcel A, Burgi E. On the Structure of the Folded Chromosomet of Escherichia coli. 21.

Xiang Y, Surovtsev IV, Chang Y, Govers SK, Parry BR, Liu J, Jacobs-Wagner C. 2021. Interconnecting solvent quality, transcription, and chromosome folding in Escherichia coli. Cell 184: 3626–3642.e14.

